# Extracellular vesicle bioactivity and potential clinical utility is determined by mesenchymal stromal cell clonal subtype

**DOI:** 10.1101/2024.09.05.609844

**Authors:** Savvas Ioannou, Alasdair G. Kay, Andrew P. Stone, Emma Rand, Samuel Elberfeld, William Bolton, Tony Larson, Rachel E. Crossland, Oksana Kehoe, David A. Mentlak, Xiao-Nong Wang, Chris MacDonald, Paul G. Genever

## Abstract

Mesenchymal stromal cells (MSCs) are a promising source of therapeutic extracellular vesicles (EVs), however it is not clear how heterogeneity within a non-clonal MSC population will affect the collective EV pool. Here we used immortalised clonal MSC lines, termed Y201 and Y202, to examine how MSC phenotype influences EV character and function.

Although morphologically similar, Y201 EVs were more abundant in EV biomarkers versus Y202 EVs, with an enhanced miRNA and proteomic content, predicted to contribute to an elaborate EV corona particularly abundant in RGD-containing proteins fibronectin and MFG-E8. We demonstrated that Y201 EVs, but not Y202 EVs, significantly increased the proliferation of articular chondrocytes and that the proliferative effect was mediated at least in part via an RGD (integrin)-FAK-ERK1/2 axis. Both Y201 and Y202 EV subsets significantly reduced proliferative index scores of activated T cells. However, only Y201 EVs, not Y202 EVs, suppressed disease activity compared to controls in different *in vivo* models of inflammatory peritonitis and arthritis.

EVs released by closely related MSC subtypes within the same heterogeneous population differ significantly in terms of cargo abundance, bioactivity, and pre-clinical *in vivo* efficacy. Analysis of defined EV subsets will aid mechanistic understanding and prioritisation for EV therapeutics.

## Introduction

Mesenchymal stromal cells (MSCs, often referred to as mesenchymal stem cells) can exhibit trilineage differentiation capacity (osteogenic, chondrogenic and adipogenic) and immuno-suppressive properties (Charbord, 2010; Najar *et al*., 2016; Wilson *et al*., 2019). Largely based on these biological functions, MSCs have been used in numerous clinical trials targeting musculoskeletal and inflammatory disorders but the range of clinical indications is very broad and success is variable (Prockop, Prockop and Bertoncello, 2014; Kabat *et al*., 2020; Rodríguez-Fuentes *et al*., 2021; Wilson *et al*., 2021; Galderisi, Peluso and Di Bernardo, 2022). Although phase 1 and 2 clinical trials continue to rise, very few reach phase 3 (Kabat *et al*., 2020; Rodríguez-Fuentes *et al*., 2021). Progress has been hampered largely by the use of non-clonal, heterogeneous and uncharacterised MSC cultures and lack of mechanistic understanding (Wilson *et al*., 2021). In addition, there is limited evidence of MSC engraftment *in vivo* (von Bahr *et al*., 2012), and disease resolution may be the direct result of MSC apoptosis (Galleu *et al*., 2017) and/or the paracrine effects of the MSC secretome, rather than the cells *per se* (Haynesworth, Baber and Caplan, 1996; Tögel *et al*., 2005; Timmers *et al*., 2007; Kusuma *et al*., 2017; Zagoura *et al*., 2019).

One of the key components of the MSC secretome are populations of extracellular vesicles (EVs). The term EV describes a heterogeneous population of secreted membrane vesicles (van Niel, D’Angelo and Raposo, 2018) that have the ability to exchange biological components between cells, thereby acting as signalling vehicles (Raposo *et al*., 1996; Ratajczak *et al*., 2006; Valadi *et al*., 2007). There are different types of EVs, frequently categorised as exosomes, microvesicles (MVs), and apoptotic bodies based on their secretory process and size (Théry, 2011; van Niel, D’Angelo and Raposo, 2018). Exosomes have a typical diameter of 30-150nm (Doyle and Wang, 2019) and are formed during the maturation of multivesicular bodies (MVBs). Biogenesis of intraluminal vesicles occurs by the inward budding of the limiting membrane of the MVB followed by scission (Piper and Katzmann, 2007; Hanson and Cashikar, 2012). Intraluminal vesicles are released as exosomes into extracellular space during the fusion of the MVB with the plasma membrane (Raposo and Stoorvogel, 2013). In contrast, MVs are formed by the outward budding of the plasma membrane (Tricarico, Clancy and D’Souza-Schorey, 2017) ranging in size from 100nm to 1000nm in diameter (Doyle and Wang, 2019). Apoptotic bodies are larger membrane-bound fragments (50-5000nm in diameter) that are generated as cells undergo apoptosis (Doyle and Wang, 2019). Apoptotic bodies are no longer considered the inert debris of dying cells, indeed they have important functions in many aspects of tissue homeostasis (Li, Liao and Tian, 2020). During EV formation, EVs are packaged with proteins, mRNAs and miRNAs and delivered to the recipient cells by various docking and uptake mechanisms to exert a biological effect (Raposo and Stoorvogel, 2013).

There is growing interest in the use of EVs as cell-free therapies. EVs may be able to offer a more cost-effective, accessible route to clinic compared to their parent cells, with improved safety profiles and amenable transport, storage and administration options. EVs have demonstrated efficacy in several preclinical animal models, including myocardial ischaemia (Lai *et al*., 2010), kidney damage (Gatti *et al*., 2011), acute lung injury (Zhu *et al*., 2014) and inflammatory arthritis (Kay *et al*., 2021) amongst others. However, in a manner similar to MSC therapy, heterogeneity within any therapeutic EV pool will hamper clinical development.

Here we used immortalised clonal MSC lines, termed Y201 and Y202, to examine how the source MSC phenotype influences EV character and function. Y201 and Y202 were isolated from the same donor and identify as “MSCs” by surface protein expression and independent transcriptomic profiling (Kay et al. 2022; James et al. 2015). Y201 MSCs lack expression of CD317 (CD317^neg^) and are model mesenchymal *stem* cells with strong trilineage differentiation and immune-suppressive properties. Y202 (CD317^pos^) MSCs have weak differentiation capacity and appear to function primarily as immune-regulatory cells (James *et al*., 2015; Kay *et al*., 2022; Stone *et al*., 2024). Our findings demonstrate that EVs released by closely related MSC subtypes within the same heterogeneous cell population differ significantly in terms of cargo abundance, bioactivity and potential for clinical efficacy, which will advance our understanding of EV biology and parent cell selection for EV therapeutics.

## Methods

### Cell culture

Y201 and Y202 MSCs were cultured using Dulbecco’s Modified Eagle’s Medium (DMEM) (ThermoFisher Scientific, Cat: 41966) supplemented with 10% Foetal Bovine Serum (FBS) and 1% Penicillin/Streptomycin (P/S) (ThermoFisher Scientific Cat: 15140). Human articular chondrocytes (AC) were isolated from primary donors following fully informed ethical consent (LREC 07/Q1105/9). ACs were cultured in DMEM-F12 (ThermoFisher Scientific Cat: 11320-033), supplemented with 10% FBS and 1% P/S.

### Preparation of EV depleted medium

DMEM supplemented with 20% FBS and 1% P/S was centrifuged at 100,000g for 18hrs at 4°C to deplete the medium of serum-derived bovine EVs. During cell expansion, medium was diluted in equal volumes with DMEM supplemented with 1% P/S. Cells were grown at 37°C in 5% CO_2_/95% atmospheric air.

### EV isolation by sequential differential ultracentrifugation

For EV isolation, 5×10^5^ cells were seeded in T175 flasks with EV depleted medium and allowed to reach 80-90% confluence. Medium was aspirated and cells were washed three times with PBS and serum-free medium was added. Two collections of conditioned medium were performed every 24hrs and stored at -70°C, at the end of the second collection cells were counted. EVs were isolated according to the Thery *et al* protocol with minor modifications, all steps were performed at 4°C (Théry *et al*., 2006). Conditioned media were centrifuged at 300g for 5mins to remove dead cells and debris followed by a second spin at 2000g for 20mins. Supernatant was transferred to Ty45i thick-walled ultracentrifuge tubes (Beckman-Coulter Cat: 355655) and centrifuged at 10,000g for 45mins in L100-XP Beckman-Coulter ultracentrifuge to pellet the “10K” EV fraction. The supernatant after the 10,000g spin was transferred to a new Ty45i thick-walled tube and centrifuged at 100,000g for 90mins to generate the “100K” EV fraction. Both 10K and 100K pellets were re-suspended in PBS by vigorous pipetting and transferred into thick-walled micro-ultracentrifuge tubes (Beckman-Coulter Cat: 357448) and centrifuged again at 10,000g and 100,000g in a Beckman-Coulter TL100 ultracentrifuge for 45 and 90mins respectively. The final 10K and 100K EV fractions were resuspended in PBS and transferred to protein LoBind tubes.

### Analysis of EVs using Nanoparticle Tracking Analysis (NTA)

EVs were analysed using the Nanosight LM14 equipped with a green laser (532nm) and 5 videos were acquired per sample. The 10K and 100K EV fractions were diluted in PBS to a concentration of 20-150 particles per frame. A script was used obtaining 5X 60s video recordings of all events for further analysis. The experimental conditions were as follows: i) Measuring time: 5X 60s, ii) Blur: Auto, iii) Detection Threshold: 4-5, iv) Blur size: Auto, iv) Number of frames: 1499.

### Transmission Electron Microscopy (TEM)

10μl of EV suspension was deposited on a 200 mesh copper grid with a formvar/carbon support film and left for 3mins to air dry. The grids were washed with three drops of dH_2_O to remove the salts from the PBS. Negative staining was attained by adding 10μl of 1% uranyl-acetate and drawing off the excess stain. The grids were left for 30min in a dry environment. EV morphology was observed by TEM using a ThermoFisher Tecnai 12 at 120kV at 18.5K and 49K magnification.

### Treatment of Y202 cells with focal adhesion kinase inhibitor

Y202 cells were seeded on a 6-well plate at a 20,000 cells/cm^2^ using DMEM supplemented with 10% FBS and 1% P/S and left to adhere for 6hr. Cells were washed thrice with PBS and serum free DMEM was added overnight. The following day, cells were treated with 10μΜ focal adhesion kinase inhibitor (FAKi) for 1hr (PF573228) and Y201 EVs were added at a 1X, 5X, and 10X concentration for 30mins. Cells were washed with PBS to remove any unbound EVs and cell lysates were created. Briefly, cells were washed with ice-cold PBS and RIPA buffer supplemented with protease inhibitors was added (Merk; Cat 11836153001). Cells were scraped and passed through a 25G needle before centrifuging at 10,000g for 20mins. The supernatant was collected and a bicinchoninic acid (BCA; ThermoFisher Scientific, Cat: 23225) assay was performed to quantify the total amount of protein.

### Western blotting

Cell lysates and EVs were separated on a 12% SDS-PAGE gel. The separated proteins were transferred onto a nitrocellulose membrane using the iBlot2 dry blotting system (20V for 1 min, 23V 4mins, 25V 2min). Membranes were blocked for an hour with 5% Bovine Serum Albumin (BSA) in PBS-0.1% Tween buffer or TBS-0.1% Tween buffer, the membranes were incubated with the following mouse monoclonal antibodies: i) Flotillin-1 (1:500; SantaCruz: 133153), ii) CD63 (1:1000; SantaCruz: 365604), iii) CD81 (1:1000; SantaCruz: 23962), iv) Alix (SantaCruz: 166952), v) GRP78 BiP (Abcam: 21685), v) Fibronectin (1:500; Abcam:2413), vi) MFG-E8 (1:200; proteintech: 67797-1-Ig) vii) ERK1/2 (1:1000; CellSignaling Technology: 4695), viii) pERK1/2 (1:1000, CellSignaling Technology: 9101) for an hour at room temperature or overnight at 4°C. After the incubation, membranes were incubated with anti-mouse horseradish peroxidase-conjugated secondary antibodies in 1:1500 dilution. The protein bands were visualised following addition of ECL PicoPlus Chemiluminescent substrate (Cat: 34577, ThermoFisher Scientific) and captured using the iBright imaging system (ThermoFisher Scientific).

### Proteomic analysis of MSC EVs

EVs isolated from 8xT175 of cells were added to 8M urea with 20mM HEPES and a phosphatase inhibitor cocktail comprising 1mM sodium orthovanadate, 1mM β-glycerophosphate and 2.5mM sodium pyrophosphate. Samples were reduced with 5mM dithiothreitol at 55°C for 30mins before alkylating with 15mM iodoacetamide for 30mins at room temperature. Solutions were diluted to 2M urea with aqueous 50mM ammonium bicarbonate before digesting with the addition of 0.5mg of sequencing grade trypsin/Lys-C protease mixture (Promega. Cat: V5071) and incubated at 37°C. Digestion was stopped after 16hr with the addition of trifluoroacetic (TFA) to 0.1% (v/v). Resulting peptides were analysed over 1-hour LC-MS acquisitions using an Orbitrap Fusion (Thermofisher). Peptides were eluted into the mass spectrometer from a 50cm C18 EN PepMap column. Three biological replicates for each cell line were run. Tandem mass spectra were searched against the human subset of the UniProt database using Mascot and peptide identifications were filtered through the Percolator algorithm to achieve a global 1% false discovery rate (FDR). Identifications were imported back into Progenesis QI and mapped onto MS1 peak areas. Peak areas were normalised to total ion intensity for all identified peptides. Relative protein quantification was performed using relative peak areas of non-conflicting peptides. Proteins were accepted for analysis provided they were detected with ≥2 peptides and ≥1 unique-peptide in at least one sample. Fold changes and p-values for differential abundance were calculated in Progenesis QI by ANOVA.

### Lipidomics analysis of Y201 and Y202 MSC EVs

Y201 and Y202 EVs were isolated by ultracentrifugation and their lipids were extracted as previously described by (Blandin *et al*., 2023). Briefly, 50μl of the EV suspension was mixed with 3μl of SPLASH Lipidomix standard: a mix deuterated internal standard. Next, 997.5μl of MeOH:CHCl_3_ (2:1, v/v) and 208μl 0.005N HCl were added, samples were vortexed followed by the addition of 332μl of CHCl_3_ and 332μl of ddH_2_O. The samples were centrifuged at 3600 g for 10 min at 4°C, the organic phase was collected and dried in a GeneVac for 30 min. The lyophilised samples were resuspended in 80μl of injection solvent, ACN:IPA (70:30) and the lipid composition was assessed using liquid chromatography with tandem mass spectrometry (LC-MS/MS). 2μl of the sample was injected for positive or negative acquisition mode using the Orbitrap Fusion Tribrid mass spectrometer (ThermoFisher Scientific, UK). The lipidomic data were analysed using the online tool MetaboAnalyst 5.0.

### Analysis of EV microRNAs (miRNAs)

RNA was extracted using the Total Exosome RNA and Protein Isolation kit (Invitrogen, Cat: 4478545). Samples were thawed and diluted with 1X PBS to 200µL total volume in an RNAse-free tube. 200µL of pre-warmed denaturing solution was then added and mixed before incubating samples on ice for 5 mins. 400µl of Acid-Phenol:Chloroform was then added to each sample and they were vortexed for 60 seconds before centrifuging at 13,000g for 5 mins at room temperature. The aqueous (upper) phase was then transferred to a fresh RNAse-free tube and the volume recovered was recorded and then used in purification. The elution solution was pre-heated to 95°C and 100% ethanol left at room temperature. The aqueous phase was diluted with 1.25 volumes of 100% ethanol and mixed. The aqueous-phase/ethanol mix was loaded onto a filter cartridge in a fresh tube and centrifuged at 10,000g for 15 seconds. The flow-through was discarded and the cartridge was then washed by centrifuging with 700µL of miRNA wash solution 1 at 10,000g for 15 seconds. Flow-through was discarded and then 2 washes with 500µL of miRNA wash solution 2/3 were performed at the same settings. The cartridge was dried with a spin of 10,000g for 1 min. The cartridge was then removed and placed into a fresh collection tube and the RNA was eluted using 50µL of pre-heated elution solution and centrifuging at 10,000g for 30 seconds. The eluate was then passed through the cartridge a second time to improve yield. After RNA purification, the samples were concentrated using Amicon Ultra 0.5ml centrifugal filters (Sigma-Aldrich, Cat: UFC500308) by first topping up to 400uL with RNAse-free water. Samples were centrifuged at 14,000g for 88 mins before inverting and collecting RNA in a fresh tube at 8000 g for 2mins. Total RNA was then quantified on a Bioanalyzer 2100 (Applied biosystems) using a Pico chip.

5-3µL of RNA was used in the NanoString miRNA ligation reaction. NanoString was performed following the manufacturer’s miRNA sample preparation protocol using the nCounter Human v3 miRNA expression assay codeset (NanoString). miRNA counts were normalised using miRNA spike-ins for other species following the NanoString procedure. miRNAs with less than 20 counts ≥1 sample were filtered out and comparisons of EV miRNAs were performed in the R programming language. The TargetScan database was used to identify predicted targets of miRNAs. Targets were included in analyses if satisfying a threshold cut-off of a context score <-0.6 (the more negative a score the more evidence that the gene is a true target of the miRNA) (Agarwal *et al*., 2015).

### Gene Ontology term enrichment and clustering

Gene Ontology (GO) enrichment was performed using the ClueGO plugin for the Cytoscape software package (Shannon *et al*., 2003; Bindea *et al*., 2009). Gene lists were assessed for enrichment against the Biological Process and Molecular function GO genesets with Benjamini-Hochberg False Discovery Rate (FDR) corrected p-values. Redundancy of GO terms was reduced by using the GO-fusion setting in ClueGO before automated clustering of significant GO terms (FDR-corrected p<0.05). Cluster diagrams were generated from ClueGO results using the AutoAnnotate plugin to facilitate organisation and labelling of like-terms and to generate titles for clusters based upon common words (Kucera *et al*., 2016). Clusters were moved to aid visualisation.

### KEGG Pathway enrichment

Lists of significantly more abundant genes and proteins were analysed for pathway enrichment against the curated Kyoto Encyclopedia of Genes and Genomes (KEGG) database using the Molecular Signatures Database website on version 7.2 (Kanehisa and Goto, 2000; Subramanian *et al*., 2005; Liberzon *et al*., 2011). Enrichment was performed for significantly different protein lists and results filtered to exclude terms with FDR corrected p-values (q) of >0.05. To minimise the effect of confounding and relatively uninformative terms, a filter excluded gene-sets containing more than 500 genes. Where p-values for enriched pathways were the same, samples were ordered by the MSigDB k/K ratio where k = the number of proteins identified in the geneset and K = the total number of proteins in that set.

### CFSE-EV labelling

Y201 and Y202 100K EVs were isolated by differential ultracentrifugation and labelled using carboxyfluorescein diacetate succinimidyl-ester (CFSE) (Invitrogen, Cat: C34554). EVs were incubated with 20μM CFSE for 120mins at 37°C in a final volume of 300μl PBS. Labelled EVs were pooled by ultracentrifugation at 100,000g for 90mins and resuspended in PBS to remove the unbound CFSE dye. Labelled EVs were transferred to a protein LoBind Eppendorf tube for storage.

### Flow cytometric determination of CFSE labelled EVs

The Cytoflex S system was used with the violet filter being set up as the primary filter to aid in the detection of small particles such as EVs. The Cytoflex S underwent a deep clean using the recommended flow cytometry detergent followed by a 20min wash with dH_2_O to remove any microbubbles in the capillary. PBS was used to calibrate background noise using the violet side scatter channel, as previously recommended (McVey, Spring and Kuebler, 2018). The acquisition settings for the Forward Side Scatter (FSC) were set at 50, FITC at 371 and Violet Side Scatter (VSSC) at 50. Gates to distinguish EVs from the background were set up using a density plot (FSC-H vs VSSC-H), and the primary threshold of the machine was adjusted to 600 to minimise the detection of buffer noise on the EV gate. The flow rate of the sample was set at 10μl/min, and events were recorded after allowing the sample to run for the first 60s to ensure a constant flow of the sample. 10,000 EVs were recorded per sample, and the median fluorescence intensity of the EVs was normalised against Y201 EVs. PBS washes were performed between each sample to ensure the removal of all EVs, avoiding cross-contamination of the sample readings.

### Analysis of EV uptake by flow cytometry

Y202 and Y201 cells were detached from plastic using 0.05% Trypsin-EDTA and resuspended in DMEM medium that was primed in advance in a 37°C, 5% CO_2_ incubator. Cells were counted using an automatic cell counter and transported in LoBind protein tubes (100,000 cells/tube). Y202 and Y201 cells were treated with Y201 and Y202 CFSE-EVs and unlabelled-EVs respectively at a 10X concentration for 1hr, 2hr, 4hr, 6hr, 8hr and 10hr.

For the RGD-blocking experiments, cells were treated with 200μΜ Gly-Arg-Gly-Asp-Ser-Pro (GRGDSP) or Gly-Arg-Ala-Asp-Ser-Pro (GRADSP) peptides (Sigma-Aldrich, Cat: SCP0157, SCP0156) for 6hrs before peptide removal by pelleting the cells and resuspending them in fresh DMEM medium. CFSE-EVs and unlabelled-EVs at a 10X concentration were introduced to cells for 4hr. Cells were pelleted by centrifugation at 300g for 5mins at 4°C and washed twice with ice-cold PBS to remove any EVs that had not been uptaken. Cells were resuspended in 200μl flow buffer (1% FBS in PBS) and EV uptake by the Y202 cells was determined by flow cytometry using the LX375 CytoFlex. The gains were adjusted as follows: i) Forward Scatter at 20, ii) Side Scatter at 40 and iii) Fluorescein isothiocyanate to detect the CFSE fluorescence at 40.

### Analysis of EV uptake by confocal microscopy

Y202 and Y201 cells were seeded on glass coverslips at a 5000 cells/cm^2^ density and left to adhere overnight at 37°C, 5% CO_2_. Y201 and Y202 CFSE-EVs or unlabelled EVs were introduced to Y202 and Y201 cells respectively at a 10X concentration for 4hr. Cells were fixed for 10 mins using 3% PFA (Thermo Scientific, Cat: 28906) followed by 3X 5 mins washes with 1mg/ml NaBH_4_ in PBS to quench free aldehyde groups followed by three washes with PBS. Cells were stained with Phalloidin CruzFluor 594 Conjugate (Santa Cruz, Cat: 363795) for 90 mins in 1% BSA in PBS. Cells were washed gently with 0.2% BSA in PBS to remove the excess Phalloidin and a mounting medium with 4’,6-diamidino-2-phenylindole (DAPI, Sigma-Aldrich, Cat: DUO82040) was used to prepare the coverslips for microscopy. Cell imaging was performed using a ZEISS LSM 980 laser-scanning confocal microscope (Carl Zeiss, Germany) with a 63X 1.4 Numerical Aperture oil objective. Solid state lasers of 405nm (blue), 488nm (green), and 561nm (red) excitation wavelengths were used to excite fluorescence with emissions collected at, 415-478nm, 491-553nm, and 561nm, respectively.

### CyQuant proliferation assay

Y202 and Y201 cells were seeded in a 96-well plate at a 9,000 cells/cm^2^ density and left to adhere overnight. The following morning, Y202 cells were treated with Y201-EVs and Y201 cells were treated with Y202 EVs at 1X, 5X and 10X concentrations. The treatments were calculated using the following equation: 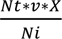 where Ni is the number of donor cells, Nt number of recipient cells, v is the volume of the EV suspension and X is the number of the desired treatment. Cells were subjected to daily EV treatments and microplates were frozen down until processing. To assess the proliferative effect of the EVs the CyQuant proliferation assay kit (Invitrogen, Cat: C7026) was used. Working CyQuant solution was prepared by diluting the cell-lysis buffer stock solution 20-fold and the CyQuant GR stock solution 400-fold using dH_2_O. Cells were incubated in the working CyQuant solution for 5-10mins. Quantification of cellular DNA per well was achieved by shaking the microplates for 2mins at 300rpm and measuring the fluorescence at 480/502nm (excitation/emission) using the CLARIOstar plate reader.

### Analysis of cell growth by ptychography

Y202 and Y201 cells were seeded in a 24-well plate at a 4,500 cells/cm^2^ density and left to adhere overnight. Y202 cells were treated with Y201 EVs and Y201 cells were treated with Y202 EVs at a 10X concentration. The microplates were loaded onto the LiveCyte microscope and cell proliferation was monitored for 72hrs by ptychographic quantitative phase imaging and analysed using the Phasefocus software v3.5. To counteract low contrast between Y202 cells and the plastic, a customised protocol on the Phasefocus analysis software was used. For example, i) rolling ball radius at 10μm and ball height at 5μm, ii) smooth image at 18μm, iii) threshold of the local maxima at 0.3, iv) consolidate seed points at 0.1, v) background and feature threshold at 0.05 and 0.5 respectively, vi) fuzzy distance Threshold and Phase weighting at 0.2μm and 6. When background particles were detected, gates were applied to remove them from the analysis, the size of the gates were depended based on the particle/cell size differences.

Articular chondrocytes (AC) isolated from donors with osteoarthritis (K257AC, K268AC, K269AC), were seeded in 24-well plates at a 3,000 cell/cm^2^ and left to adhere overnight. Cells were treated with Y201 100K EVs at a 10X and 20X concentrations and cell proliferation was monitored for 72hrs on a LiveCyte microscope as above. ACs were also treated with 200μΜ GRGDSP or GRADSP peptides (Sigma-Aldrich, SCP0157, SCP0156) for 6hrs. Cells were washed twice with PBS and Y201 100K EVs were added at a 20X treatment. Proliferation rate was monitored for 72hrs using the LiveCyte microscope.

### Scratch wound assay

Y202 cells were seeded in 24-well plates at 30,000 cells/cm^2^ and left to adhere for 6-8hrs. Cells were washed with PBS and cells were serum-starved overnight. A 10μl pipette tip was used to create a scratch in the confluent monolayer of cells and washed with PBS to remove dead cells and debris. Y202 cells were treated with 10X concentration of 100K EV fraction from Y201 cells and wound closure was monitored over 24hrs by ptychographic quantitative phase imaging with the LiveCyte microscope. Images were analysed using the phasefocus software v3.5 Single cell analysis was performed using the MTrackJ plugin in ImageJ and measurements were imported into the chemotaxis tool to calculate the cell velocity, total track length, Euclidean distance, directness and forward migration index.

### Chondrogenic differentiation assay

Primary MSCs were isolated from femoral head donations from three osteoarthritis patients following ethical approval (LREC 07/Q1105/9). Prior to formation of chondrogenic micromass pellets, primary MSCs were exposed for 6hrs to 10X 100K EVs (EVs derived from 10X Y201 or Y202 MSCs). Following this, micromass pellets were formed at a density of 2.35×10^5^ cells per pellet in 100μl growth media in microcentrifuge tubes by centrifuging at 300g for 5mins and then incubating overnight at 37°C.

Pellets were detached from the tube and appropriate medium added up to 1.2ml taking care not to disrupt the pellet. Negative controls received basal medium comprising DMEM containing sodium pyruvate and L-glutamine, supplemented with 1% Insulin-Transferrin-Selenium (ITS)+3 (Sigma-Aldrich, Cat: I2771), 40µg/ml L-Proline (Life Technologies, Cat: P6698), and 1% non-essential amino acids (Sigma-Aldrich, Cat: TMS-001-C). Chondrogenic medium was composed of basal medium with the addition of 0.1µM dexamethasone (Sigma-Aldrich, Cat: D2915), 50µg/ml L-Ascorbic Acid (Sigma-Aldrich, Cat: A4544) and 10ng/ml TGF-ß1 (Peprotech, Cat: 100-21). Tubes were incubated at 37°C and medium changed every 3-4 days. Test conditions were treated once per week with 10X EVs per pellet. At day 7, pellets were removed and rinsed twice with 1ml PBS then fixed for 10mins with cold paraformaldehyde. Pellets were again washed twice with PBS and paraffin wax embedded within 48hrs of fixation using a Leica tissue processor on the integrated ‘small biopsy’ program. Pellets were then sectioned at 6μm thickness. To ensure consistency, pellets were sectioned to a depth of 48 microns and then 8 further sections of thickness 6μm collected for testing. Paraffin embedded sections were cleared and rehydrated prior to staining with 0.02% Fast Green for 5mins and 0.1% Safranin O for a 15-minute period. Following staining, sections were dehydrated and mounted using DPX mounting medium prior to imaging at 20X resolution with Z-stack imaging on an Axio Scan.Z1 slide scanner.

### T cell activation assay

To determine MSC-derived EV immunomodulation for deactivation and suppression of T cell proliferation, suspension cultures of 1.0×10^5^ primary human peripheral blood-derived CD4+ T cells (Stem Cell Technologies) were grown in RPMI-1640 with 10% FBS 1% P/S (Gibco, Cat: 10363083) and were pre-treated for 6hrs with 20X EVs isolated from serum-free conditioned medium collected over 24hrs culture from 2.0×10^6^ of Y201 or Y202 MSCs. For positive controls, 1.0×10^4^ MSCs were seeded into a 96-well U-bottomed plate and cultured for 24hrs at 37°C, 5% CO_2_ prior to addition of 10X peripheral blood-derived CD4+ T cells. Continual proliferative capacity was assessed as a measure of T cell proliferation. CD4+ T cells were stained for 10mins at 37°C using 1μM VPD450 Violet proliferation dye (eBioscience, Inc., Cat: 65-0863-14). T cells were activated using anti-CD3ε/CD28 Dynabeads (Gibco, Cat: 10310614) at a bead-to-cell ratio of 1:1 then seeded at a density of 1.0×10^5^/well (ratio 10:1) in 200μl RPMI-1640 with 10% FBS, 0.05μg/mL IL-2 (Peprotech, Inc, Cat: 200-02). T cells seeded alone (no treatment) or seeded onto 1.0×10^4^ Y201 or Y202 MSCs were applied as negative and positive controls respectively and all conditions were tested with and without activation. Plates were cultured for 6 days at 37°C prior to removal of Dynabeads with the DynaMag-2 as per manufacturer’s recommendations. T cell proliferation was assessed with flow cytometry, with reduction in signal intensity visualised for peaks using FCS Express 7.0 proliferation analysis. Proliferation was assessed through VPD450 dilution (diminished staining intensity) described through a proliferative index (PI) calculated from the fluorescence intensity at each cell division as described previously (Kay *et al*., 2022). Proliferative cycles undertaken were calculated on 50% fluorescence intensity reduction peaks, measuring from fluorescence intensity at Day 0 to the final division detected.

For assessment of T helper differentiation, T cells were activated and cultured with MSC-derived EVs (Y201 and Y202) or with MSC monolayers, as described above excluding VPD450 staining. Following 6 days of culture, T cells were re-stimulated using a combination of phorbol 12-myristate 13-acetate (PMA) (50ng/ml) (Sigma-Aldrich, Cat: P8139) and Ionomycin (1μg/ml) (Invitrogen, Cat: I24222) and intracellular cytokines retained using transport inhibitor cocktail with 10μg/ml brefeldin A and 2μM Monensin (Invitrogen, Cat: 00-4980-93). T cells were cultured for 4hrs at 37°C then stained for surface marker CD4 (FITC) (Life Technologies, Cat 11-0049-42) prior to fixation and permeabilisation for intracellular staining of anti-human IFN-γ (Th1) (PE), IL-4 (Th2) (APC) or IL17a (Th17) (PE-Cy7) (Life Technologies, Cat: 12-7319-82, 17-7049-42, 25-7179-42); or CD4 and CD25 (Life Technologies, Cat: 17-0257-42) then fixation/permeabilisation and staining for nuclear protein FOXP3 for regulatory T cells (Life Technologies, Cat: 12-4776-42). All cells were measured using the Cytoflex LX flow cytometer and analysed with FCS Express 7. Comparisons were drawn for percentage of T helper differentiation within the CD4+ cell population and signal intensity (Median) for each antibody tested.

### *In vivo* assessment of immunomodulatory capacity in a murine peritonitis model

An *in vivo* peritonitis model was used in C57BL/6J mice aged 8-10 weeks with zymosan and schistosome egg irritant for induction of inflammation. These experiments were carried out in accordance with the Animals and Scientific Procedures Act 1986, under UK Home Office Licence (project licence number PPL PFB579996 approved by the University of York Animal Welfare and Ethics Review Board). At day 0, mice were administered with an intraperitoneal infusion of 1mg of zymosan A (Merck) or 5000 schistosome eggs in 200μl of PBS. Immediately following administration of irritant, test condition mice were administered an intraperitoneal infusion of EVs isolated from serum-free conditioned media collected over 24hrs from either 4.0×10^7^ of Y201 MSCs in 100μl of PBS to treat zymosan-induced inflammation or 2.0 x 10^7^ of Y201 in 100 μl of PBS to treat schistosome eggs-induced inflammation; negative control mice were given PBS vehicle only.

After 24hrs, mice were euthanised using CO_2_ overdose and cervical dislocation. Intraperitoneal injection of 4ml of ice cold RPMI-1640 was administered as peritoneal lavage. The process was repeated with a second 4ml RPMI-1640 wash and solutions pooled to form the peritoneal exudate cells (PEC). For each animal tested, red blood cells were lysed from the PEC using Red Cell Lysis buffer (Merck) and a cell count performed. PEC samples were initially stained for Ly6C (APC), F4/80 (PE-Cy7), CD45 (PerCP-Cy5.5) (Biolegend, Cat: 128016, 123114, 103132) and Ly6G (FITC), CD11b (BUV395) and SiglecF (BV421) (BD, Cat: 551460, 563553, 562681). PEC samples were then stained for TCRb (AF488), CD3 (APC-Cy7), CD4 (PerCP-Cy5.5), CD62L (APC) and CD44 (PE) (BioLegend, Cat: 109215, 100222, 100540, 104412, 103008). For all tests, Zombie Aqua (BioLegend, Cat: 423101) was used to exclude dead cells.

### Adjuvant-Induced Arthritis Model

The adjuvant-induced arthritis (AIA) model of inflammatory arthritis was used in male C57Bl/6 mice aged 7–8 weeks as previously described (Kehoe *et al*., 2014) with all procedures performed under Home Office project licence PPL40/3594. Treatments were administered as intra-articular injections of EV suspensions in 15μL of PBS corresponding to the number of EVs produced by ∼5.0×10^6^ cells, or PBS only controls. Mice were injected 1 day post arthritis induction using 0.5 ml monoject (29G) insulin syringes (BD Micro-Fine, Franklyn Lakes, NJ, USA) through the patellar ligament into the knee joint. Swelling was determined by measuring knee joint diameter at 1-, 2- and 3-days post-injection using a digital micrometre (Kroeplin GmbH, Schlüchtern, Germany). Peak swelling caused by inflammation was observed at 24hrs post-induction and diameters are reported as reduction from peak swelling. Four independent experiments were performed to determine the effect of EVs on disease activity. Animals were sacrificed for histological analysis at day 3 post arthritis induction. Joints were fixed in 10% neutral buffered formal saline and decalcified in formic acid for 4 days at 4 °C before embedding in paraffin wax. Tissue sections (5μm) were stained with haematoxylin and eosin (Merck Life Science UK Ltd., Cat: HHS32, HT110132) and mounted in Hydromount (Scientific Laboratory Supplies Ltd., Cat: Nat1324). H&E sections were scored for synovial hyperplasia (0 = normal to 3 = severe), cellular exudate (0 = normal to 3 = severe) and synovial infiltrate (0 = normal to 5 = severe) by two independent observers blinded to experimental groups. The scores were summated to produce a mean arthritis index.

### Statistical analysis

The statistical analyses for all experiments were performed using GraphPad Prism v9.0.2. The statistical significance between treatment and control (DMEM) was assessed by *t*-test, One-Way ANOVA or Two-Way ANOVA and Bonferroni or Tukey corrections. Error bars show Standard Error of the Mean (SEM) and asterisks represent the following P values, *p<0.05, **p<0.01, ***p<0.001, ****p<0.0001.

## Results

### Characterisation of MSC EV subtypes

The size and yield of EVs derived from Y201 and Y202 MSCs were determined by Nanoparticle Tracking Analysis (NTA). We identified three populations in the Y201 10K fraction with peak sizes of 100 ± 18.25nm, 112 ± 3.8nm, and 211 ± 12.6nm. The Y202 10K fraction had three distinct populations with peak sizes of 96 ± 8.6nm, 130 ± 0.5nm and 171 ± 4.2nm (Figure 1A). The mean size of vesicles isolated from the Y201 and Y202 100K fractions was calculated to be 133.5nm and 124.8nm respectively. In the Y201 100K fraction we identified one distinct population with a modal size of 104 ± 12nm and a second peak at 112 ± 1nm. The Y202 100K fraction also had one distinct population with a modal size at 99 ± 8nm and another peak at 114 ± 2.2nm (Figure 1A). The mean size of vesicles isolated from the Y201 and Y202 100K fractions was calculated to be 133.5nm and 124.8nm respectively. Collecting EVs from non-proliferating cells allowed the estimation of the secreted EVs per million cells. There were no significant differences between Y201 and Y202 10K EV yields, whereas Y201 cells secrete a significantly higher number of EVs isolated in the 100K fraction compared to Y202 100K EVs (Figure 1B). Both the 10K and 100K fraction of Y201 and Y202 EVs had typical morphological characteristics, as demonstrated by negative staining followed by TEM. The TEM images were analysed with the TEMExosome Analyser software for morphometric quantification of the EVs (Kotrbová *et al*., 2019). In general terms, the Y201 10K EVs were more variable and larger than the other fractions but there were no significant differences between Y201 and Y202 100K EVs in any morphological parameters measured (Figure 1C-D). TEM analysis showed the Y201 100K fraction had a mean diameter of 138.5nm whereas the Y202 100K fraction had a mean diameter of 116.3nm, corresponding closely with the NTA data. Western blotting demonstrated that Flotillin-1, Alix, CD63, CD81 were enriched in the Y201 EV fractions, particularly in 100K, compared to Y202 EV fractions (Figure 1E). The presence of the endoplasmic reticulum protein, binding immunoglobulin protein (BiP), in the cell lysates and absence in the EV fractions confirmed the purity of the EV suspensions. Considering these data, subsequent work focused on the 100K EV fraction.

**Figure 1.**
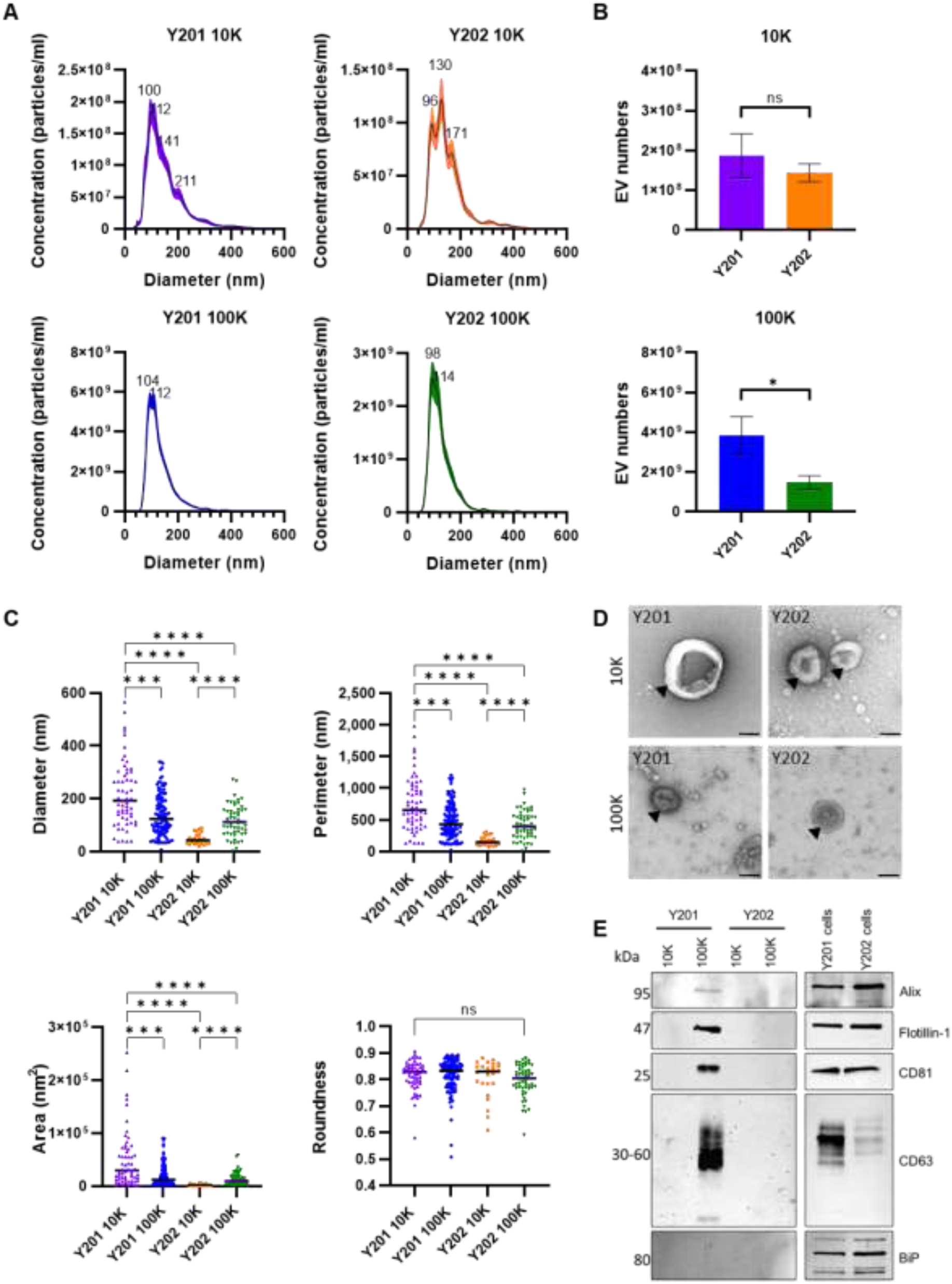
Characterisation of Evs isolated from MSC subtypes. A) Nanoparticle tracking analysis was used to determine sizes of the 10K and 100K EV fractions from Y201 and Y202. B) Quantification of EVs using the LM14 Nanosight 3.4 software. C) Morphological analysis of TEM images was performed using TEM_ExosomeAnalyzer to determine diameter, perimeter, area and roundness. D) Negatively stained EV samples imaged using TEM, Scale bar: 100nm. E) Western blot analysis of common EV markers (Alix, Flotillin-1, CD81, CD63) and negative control (BiP). Tukey’s (One-Way ANOVA) statistical test *p<0.05,,***p<0.001, ****p<0.0001.

### Analysis of Y201 and Y202 miRNA cargo

We next analysed the miRNA content of EVs (100K fraction) from Y201 and Y202 cells. miRNA detection was achieved using the Nanostring nCounter using a codeset with 828 miRNA probes; miRNAs with more than 20 copies on average were considered present in the 100K EV fraction. Y201 cells packaged 33 miRNAs, whereas Y202 cells packaged 26 miRNAs, 19 of which are mutual for Y201 and Y202 EV populations (Figure 2A). These miRNAs were assigned to their experimentally validated target genes on the miRTarBase online database. The 33 Y201 EV-miRNAs target 673 different genes, and the 26 Y202 EV miRNAs target 1089 genes; 461 of these genes are targeted by both Y201 and Y202 EV-miRNA (Figure 2B). The fold change of the EV miRNAs calculated for all miRNAs that had, on average, more than 20 counts in either Y201 or Y202 EVs are provided in Supplementary Table S1. An unpaired *t-test* with multiple corrections using the absolute miRNA numbers from both Y201 and Y202 EVs was performed. Ten miRNAs were significantly upregulated in Y201 EVs versus Y202 EVs, and two were significantly upregulated in Y202 EVs versus Y201 EVs (Figure 2C). The largest fold change in Y201 EVs was found in miR-100-5p whilst in Y202 EVs it was miR-6721-5p. Two of the three miRNAs comprising the highly conserved miR-29 family (miR-29a-3p and miR-29b-3p) were increased in Y201 EVs versus Y202 EVs and of the miRNAs elevated in Y201 EVs, miR-125b-5p had the highest mean miRNA count. We used the TargetScan miRNA prediction algorithm to identify possible targets for the enriched miRNAs in both Y201 and Y202 EVs. The 10 miRNAs elevated in Y201 EVs produced 378 predicted targets while the two miRNAs increased in Y202 EVs targeted a possible 743 mRNAs (total-context++ -<0.6) (Agarwal *et al*., 2015). The list of potential targets for each miRNA was collectively assessed using the KEGG pathway databases. Enrichment was identified for the “Focal adhesion” and “ECM-receptor interaction” pathways for Y201 EV miRNAs (Figure 2D). Despite having a greater number of predicted targets, the Y202 EV miRNA targets did not collectively enrich for any GO terms. However, when compared against KEGG pathways, six pathways were enriched at low significance, with the “Cell adhesion molecules’’ pathway most significant (Figure 2D).

**Figure 2.**
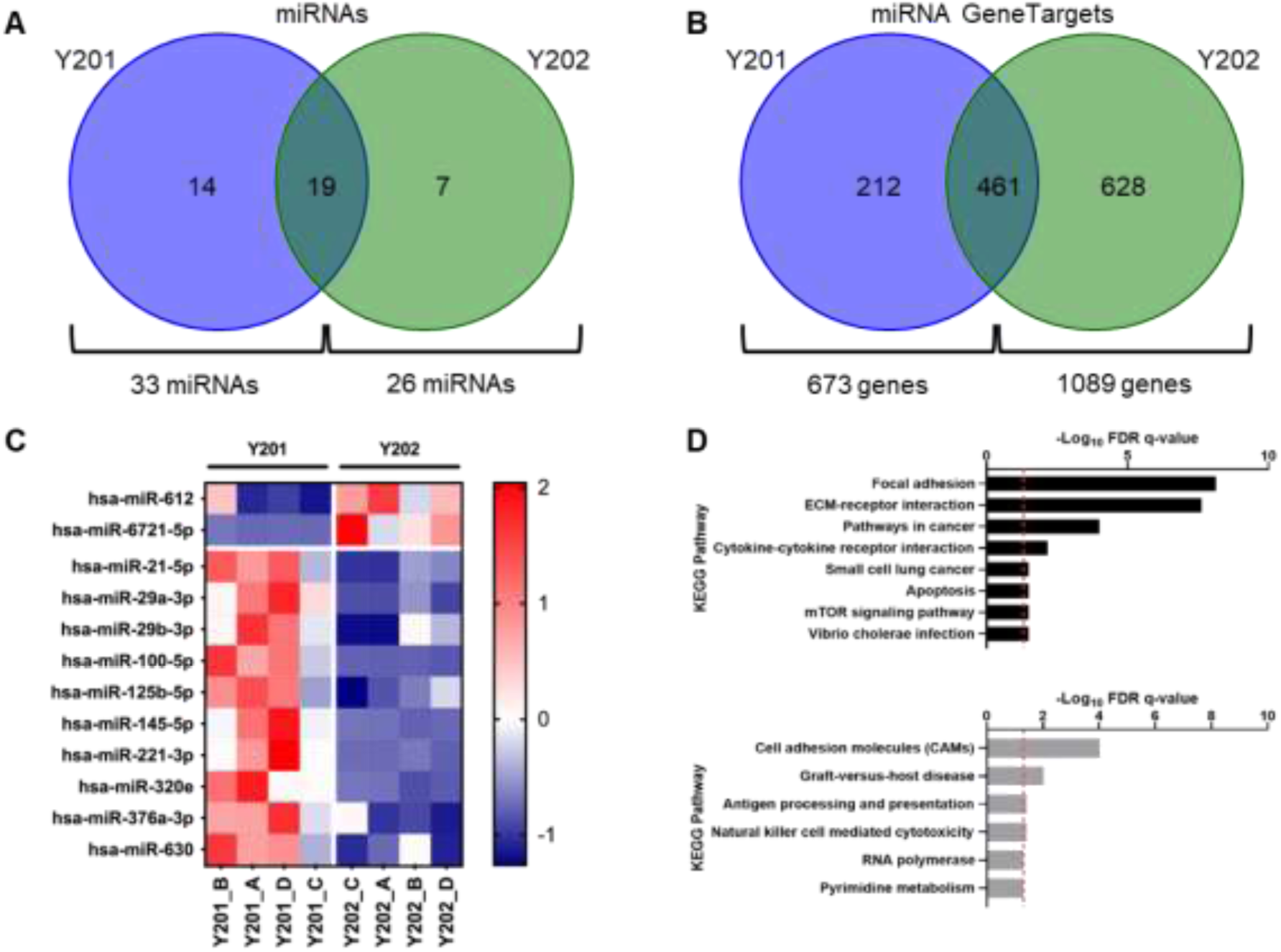
Analysis of Y201 and Y202 EV miRNA cargo. A) Presence and absence of miRNAs across Y201 and Y202 EVs displayed as Venn diagram. B) MIENTURNET miRTarBase analysis of genes targeted by miRNAs packed into Y201 and Y202 EVs miRNAs. C) Clustered heatmap of 12 significantly differently expressed miRNAs identified in EVs after Nanostring miRNA quantification between Y201 and Y202 cells. 10 miRNAs were upregulated in Y201 EVs versus 2 in Y202 EVs, n=4, ANOVA, p<0.05. Colour scale is representative of z-score. D) KEGG pathway enrichment for predicted targets of miRNAs that were more abundant in Y201 vs Y202 EVs. Significantly enriched KEGG pathways (FDR corrected q-value < 0.05) for genes that were predicted targets of the 10 miRNAs that were more abundant in Y201 EVs (black, top) and Y202 EVs (grey, bottom). Red line marks q < 0.05. q = FDR corrected p-value.

### Analysis of Y201 and Y202 EV lipid composition

LC-MS/MS was used to determine the lipid profile of EVs derived from Y201 and Y202 MSCs. We identified 172 different lipid species, though only 7 had significantly different abundances between the 2 groups with >2-fold change (p<0.05, t-test). PCA plots showed some separation between Y201 and Y202 EV datasets, though with significant overlap, which was replicated using a hierarchical clustering heatmap, indicating similar lipidomic features across the two EV populations (Supplementary Figure S1A-C).

Comparable lipid signatures for Y201 and Y202 EVs were also revealed based on the most abundant lipids. EVs derived from both Y201 and Y202 MSCs, had a high abundances of phosphatidylcholines (PCs) and sphingomyelins (SMs), particularly, PC(16:0_18:1), and SM(d18:1/16:0) (Supplementary Tables 2 and 3). We attempted to define those lipids which were significantly different in Y201-derived EVs in comparison to Y202-derived EVs, but few of these were identifiable. It is notable however, that there was a significant enrichment of the lactosylceramide LacCer(d18:1/22:0) in Y201 EVs compared to Y202 EVs (Supplementary Table 4).

### Analysis of Y201 and Y202 EV protein cargo

Next, EVs (100K) isolated from Y201 and Y202 cells were analysed by LC-MS/MS to determine protein content. Hierarchical clustering analysis was performed to compare overall abundances of shared proteins between Y201 and Y202 EVs (Figure 3A). Although both EV subtypes are relatively homogenous in their composition of proteins, there are variations in protein abundances. From a total of 663 proteins identified across all samples, there was a significantly increased abundance of 162 proteins in Y201 EVs versus 14 proteins significantly more abundant in Y202 EVs (ANOVA, p<0.05) (Figure: 3B). To investigate differential enrichment further, the top and bottom 25 most abundant EV proteins were visualised in a heatmap (Supplementary Figure S2), demonstrating that most of the proteins enriched in Y201 EVs have a larger fold change compared to those in Y202 EVs. To explore potential functional differences between the most abundant Y201 or Y202 EV proteins, enrichment analysis was conducted against the GO Biological Process geneset. In terms of Y201 enriched EV proteins, extracellular matrix (ECM) organisation and terms relating to neutrophil mediated immunity were among the most upregulated processes (Figure 3C). Statistically more abundant proteins in Y202 EVs were enriched for pathways including supramolecular fibre organisation, amyloid fibril formation, and anti-fungal humoral response (Figure 3D).

**Figure 3.**
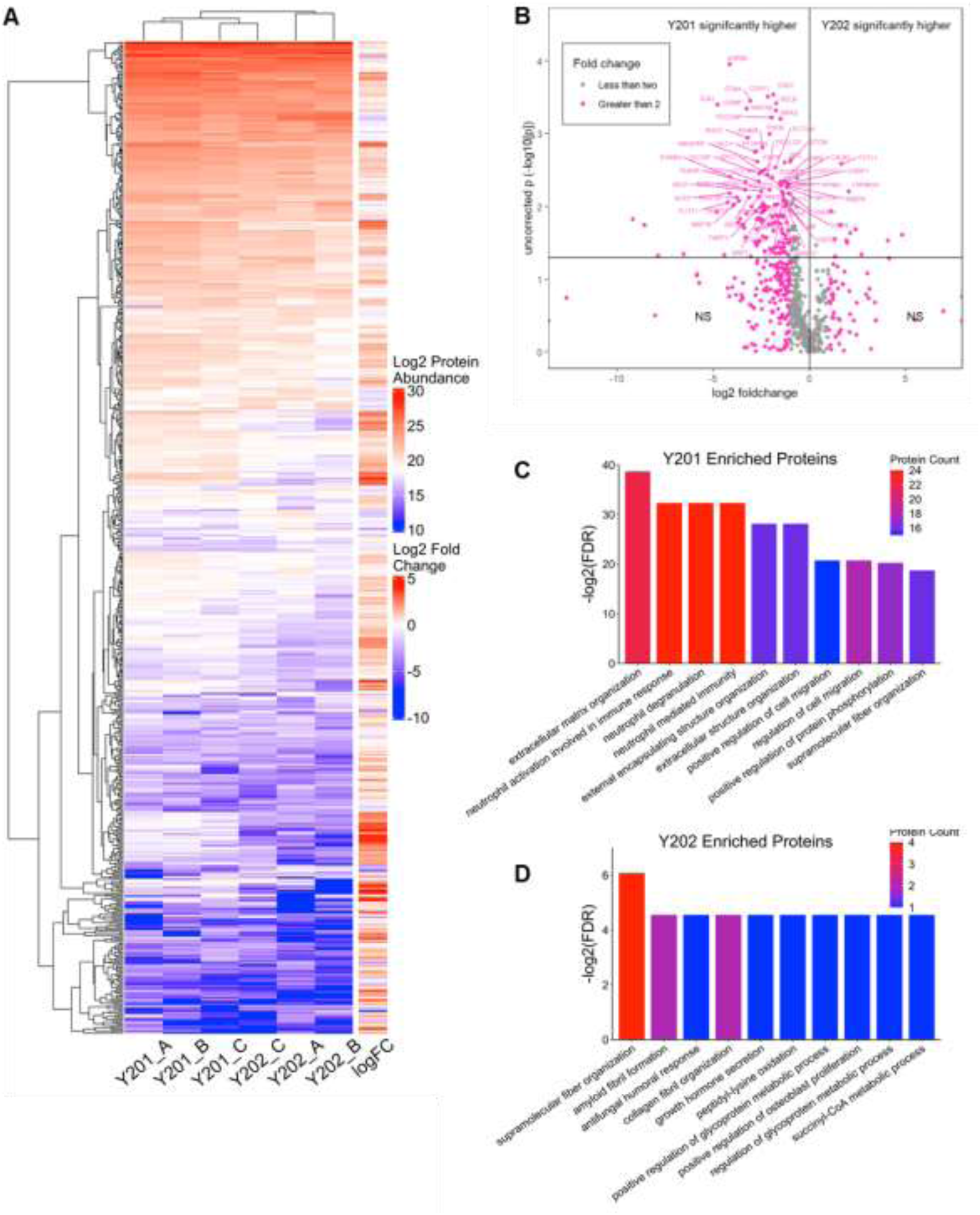
Analysis of Y201 and Y202 EV protein cargo. A) Hierarchical clustering heatmap of protein abundance for shared proteins among Y201 and Y202 EVs. Higher levels of protein abundance are indicated in shades of red, and lower levels are indicated in shades of blue. B) Volcano plot of proteins identified in EV isolations from Y201 and Y202 clonal lines. Volcano plot of fold change for the 663 proteins identified from LC-MS/MS analysis of EVs isolated from Y201 and Y202. Proteins with a fold change >2 between cell lines are shown in pink with fold changes <2 in grey. The two upper quadrants contain proteins that were significantly different between Y201 and Y202 as determined by ANOVA (p<0.05). C) Enrichment analysis against Biological Process Gene Ontology geneset for proteins enriched in Y201 EVs and D) those enriched in Y202 EVs. Shown are the top 10 most significant processes as measured by -log10 of the adjusted p-value (FDR) and total number of proteins in the dataset which match each term. Higher protein count indicated in red and lower protein count indicated in blue.

To provide further insight into the nature of the enhanced Y201 EVome, GO analysis against Biological Process, Molecular Function, and Cellular Component genesets was carried out. In terms of Biological Process, the Y201 EVome is predominantly enriched in organisation and structure of the extracellular matrix, immunomodulatory activity mediated through neutrophils, and various adhesion and migration processes (Figure 4A). The Molecular Functions of these proteins largely pertain to the binding of cadherin and various nucleic acids including GDP, nucleic triphosphates, and RNAs (Figure 4B). Enrichment against the Cellular Components geneset revealed that these proteins are significantly present in focal adhesions and cell-substrate junctions, to a relatively substantial degree. (Figure 4C). To gain a broader perspective of potential Y201 EV functions *in vivo*, the Y201 EVome was assessed for enrichment against the GO Biological Process geneset in the BiNGO plugin for Cytoscape, which formulates clustered networks of significantly enriched processes (Figure 4D). Sizeable clusters formed around similar significant processes relating to the immune system, regulation of cellular processes, developmental processes, and organisation of cellular components.

**Figure 4.**
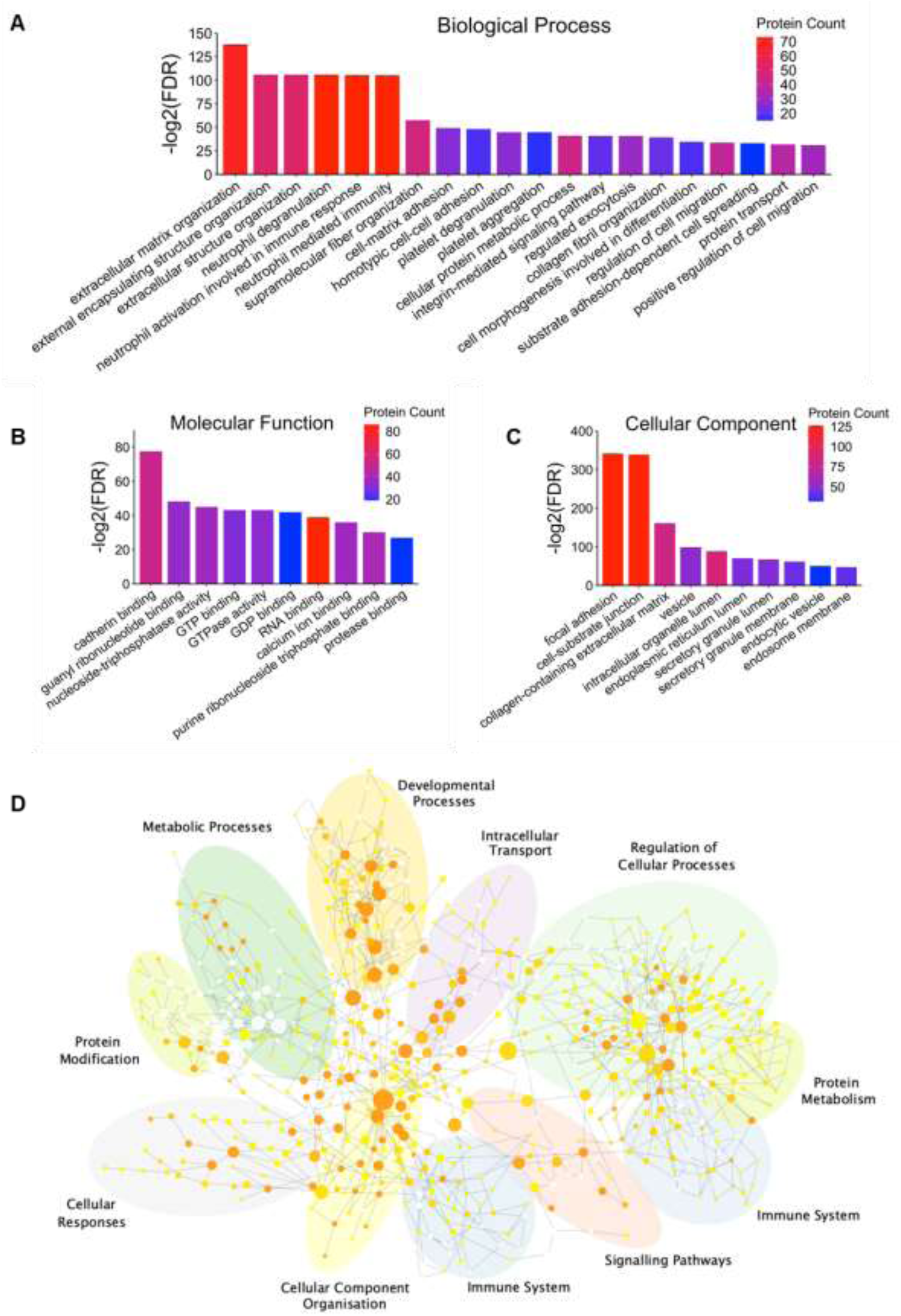
Functional enrichment analysis of the Y201 EV proteome. Enrichment analysis against A) Biological Process, B) Molecular Function and C) Cellular Component Gene Ontology (GO) terms for the Y201 proteome. Only the most significant terms are shown, as determined by -log10 adjusted p-value (FDR) and number of proteins associated with each term. Higher protein count indicated in red and lower protein count indicated in blue. (D) BiNGO network of significant GO Biological Process terms for the Y201 proteome, annotated to cluster common terms by similar processes. Darker shades of orange correspond to higher significance (p-value threshold < 0.05).

Considering the most abundant Y201 EV proteins were associated with ECM organisation, we attempted to model the extravesicular corona by constructing a protein-protein interaction (PPI) network in STRING (Supplementary Figure S3A). We identified a large number of PPIs between ECM component and cognate vesicle membrane integrins and tetraspanins. Fibronectin (FN1) and milk fat globule epidermal growth factor 8 (MFGE8) were amongst the most abundant presumptive Y201 EV coronal proteins, which we confirmed by western blot analysis relative to Y202 EVs (Supplementary Figure S3B). Collectively, these data demonstrated that EVs released by MSC subtypes isolated from the same donor, have broadly similar morphological characteristics but differ significantly in terms of yield, EV marker abundance and cargo.

### Uptake of EVs by different MSC subtypes

To determine EV uptake by recipient Y202 and Y201 cells, EVs from each cell line were fluorescently-labelled using CFSE and internalised signal measured by flow cytometry and confocal microscopy. The CFSE intensity was normalised to Y201 100K EVs, and it was demonstrated that Y202 100K EVs have significantly lower fluorescent intensity by approximately 20%. All the median values obtained using Y202 EVs were normalised against Y201 EVs for the subsequent experiments (Supplementary Figure S4). Y201 and Y202 cells were treated with Y202 and Y201 CFSE-labelled EVs, respectively, and flow cytometry was used to quantify EV uptake by the cells. A gating strategy using the Cytoflex LX375 was set up using the following parameters. An FSC against SSC plot was created to discriminate between cells and debris in suspension. After gating out the debris, single cells were distinguished by plotting FSC-H against FSC-A. Cells that were not treated with stained EVs were used to identify the autofluorescence of the cells by plotting SSC against the CFSE signal. Finally, cells treated with CFSE-EVs were used to measure the successful detection of EV uptake. An approximate signal of 3,000a.u. on the CFSE parameter was considered the background signal of both Y201 and Y202 cells (Supplementary Figure S5).

Cells were treated with unlabelled or CFSE-labelled EVs for up to 10 hrs and the extent of EV uptake was determined in living cells by flow cytometry. Y202 cells exposed to CFSE-Y201 EVs increased in fluorescence over the first 4hrs and levels remained elevated for up to 10hrs (Figure 5A, B). Orthogonal projections by confocal microscopy after 4hrs of CFSE-Y201 EV exposure, confirmed the fluorescent signal observed was from intracellular EVs in Y202 cells (Figure 5C). When CFSE-labelled Y202 EVs were added to Y201 cells, only a slight increase in fluorescence was detected after 1hr, with levels decreasing over the remaining time course (Figure 5D, E). In addition, Y202 CFSE-EVs could not be observed by confocal microscopy in Y201 cells after 4hrs (Figure 5F). These findings demonstrate that different MSC subtypes, Y201 and Y202, and the EVs they produce display distinct uptake kinetics, which is likely to impact biological function.

**Figure 5.**
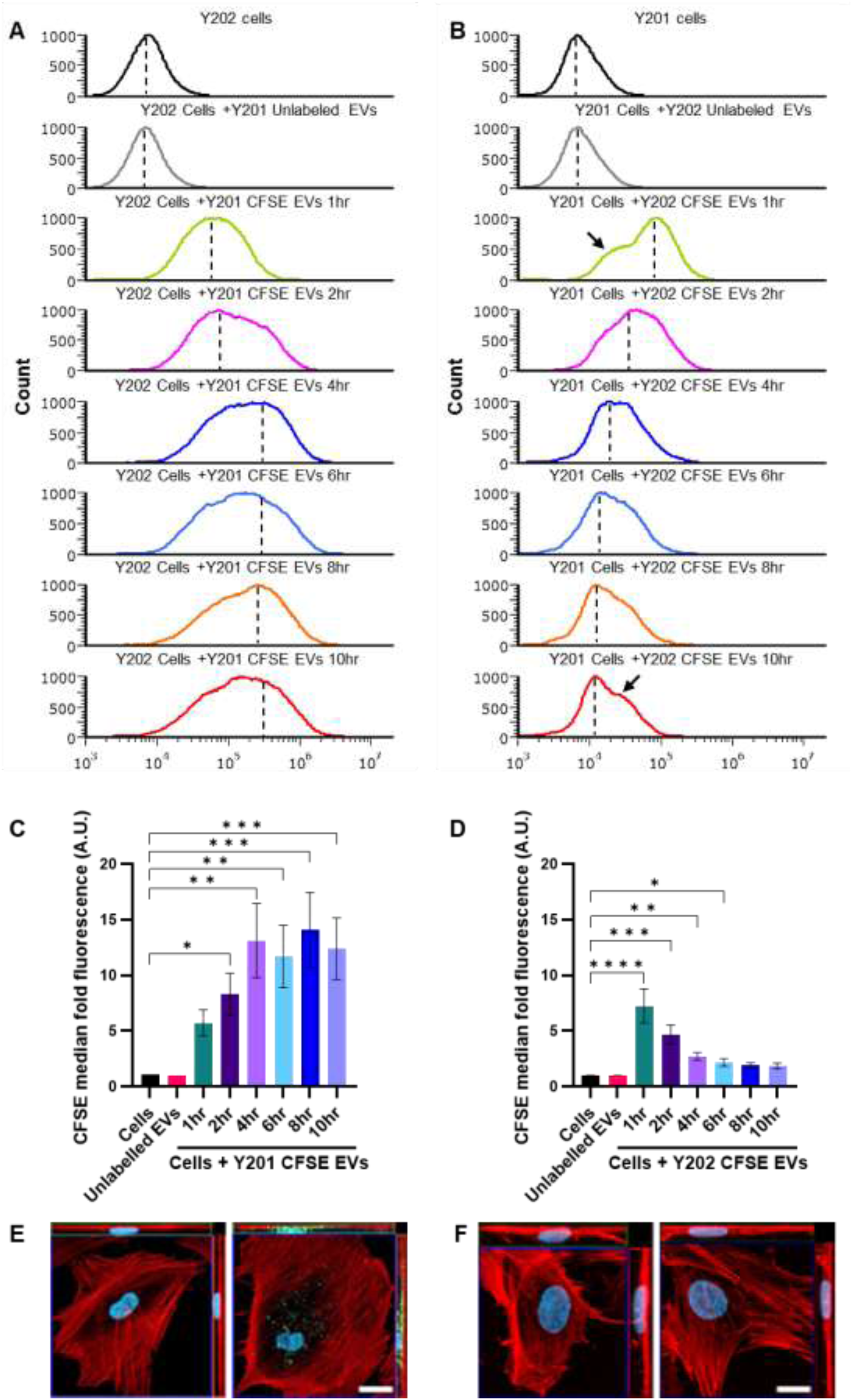
Uptake of Y201 EVs by Y202 cells (A, C, E) and Y202 EVs by Y201 cells (B, D, F). (A, B) Y202 and Y201 cells were treated with CFSE-labelled Y201 and Y202 Evs respectively. CFSE signal in the cells was monitored for 10hrs and histograms (counts vs CFSE) signals were normalised to peak values. Dashed lines represent the median values used to quantify CFSE signal levels relative to control (cells), and arrows show the presence of a second population of cells, (C, D) Quantification of the median CFSE intensity relative to control were plotted as bar graphs. n=3, error bars= SEM, Kruskal-Wallis test, *p<0.05, **p<0.01, ***p<0.001, ****p<0.0001. (E) confocal microscopy orthogonal projections of Y202 cells exposed to CFSE-labelled Y201 EVs for 4hrs and (F) Y201 cells exposed to CFSE-labelled Y202 EVs for 4hrs. Scale bar: 20μm, red= rhodamine-phalloidin, blue = DAPI, green = CFSE.

### Y201 EVs promote proliferation of Y202 cells

To test bioactivity, Y201 EVs were introduced to Y202 cells daily and cell numbers were quantified using a CyQuant assay. The 100K EV fraction from Y201 cells at 1X, 5X and 10X treatments induced a significant increase in DNA content with the 10X concentration maintaining significance 72hr after the initial treatment (Figure 6A). The proliferative effect of Y201 EVs on Y202 cells was also examined using quantitative live imaging (Livecyte ptychography microscopy) that enables simultaneous measurement of several different parameters associated with proliferation at a single cell level, including: doubling times, dry mass, cell counts and confluency. We also demonstrated that a single treatment with Y201 EVs stimulated growth of Y202 cells with doubling times significantly reduced when measured by cell count or dry mass (Figure 6B, C) with significant increases in total cell counts, total dry mass and confluency over 72hrs compared to untreated controls (Figure 6D-F). Conversely, EVs derived from Y202 cells had no significant effect on Y201 cells in the same proliferation assays (CyQuant, livecyte doubling times, dry mass, cell counts, confluency) at any time point (Figure 7).

**Figure 6.**
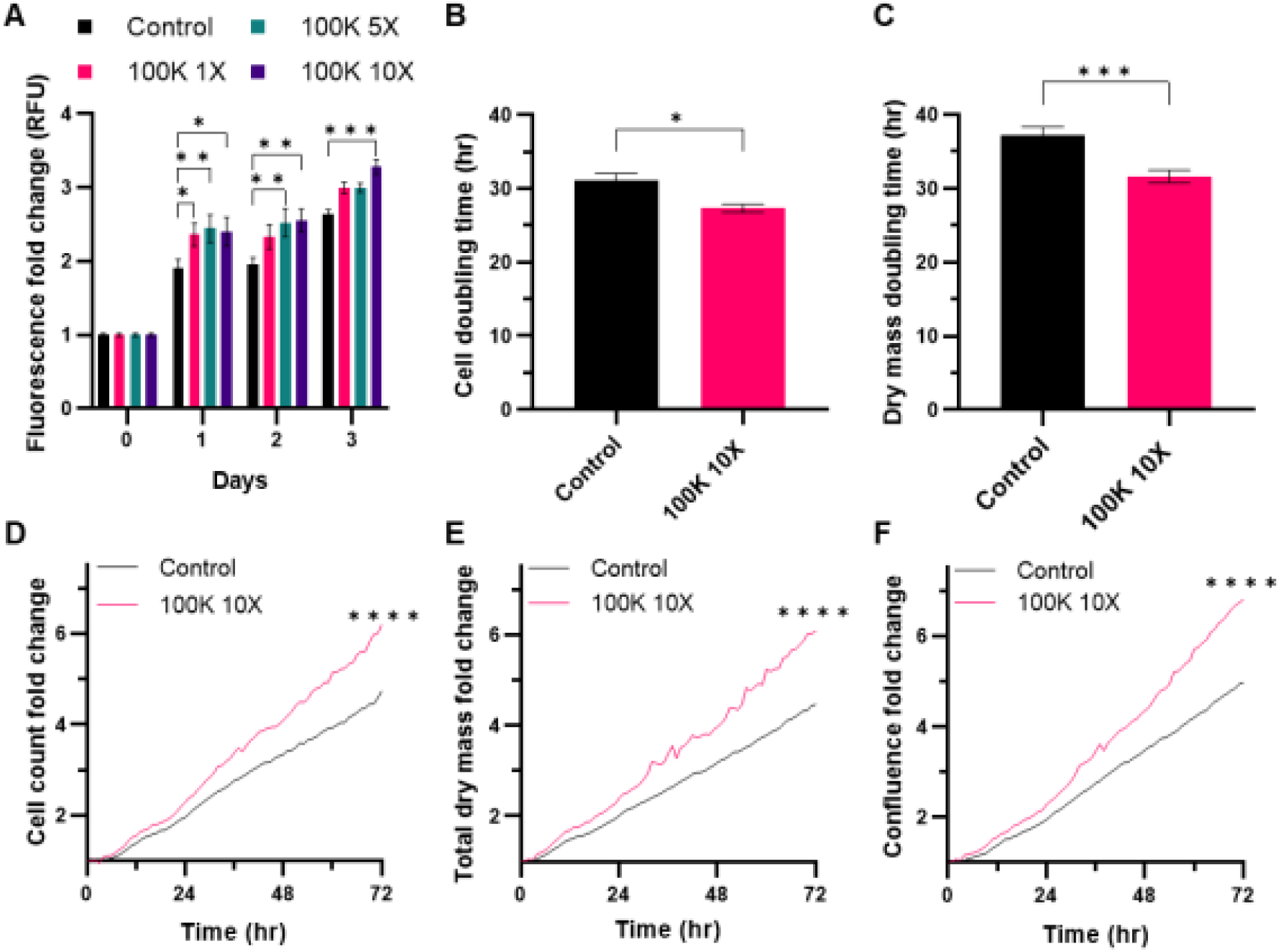
Effect of Y201 EVs on Y202 proliferation. A) Y202 cells treated with 1X, 5X and 10X concentrations of Y201 EVs and cell number determined over 72hrs (CyQuant assay). Livecyte image analysis was used to determine the effect of 10X Y201 EVs on Y202 B) cell doubling time, C) dry mass doubling time, D) cell counts, E) dry mass, and F) Y202 confluence. n=3 t-test or Two-Way ANOVA *p<0.05, ***p<0.001, ****p<0.0001.

**Figure 7.**
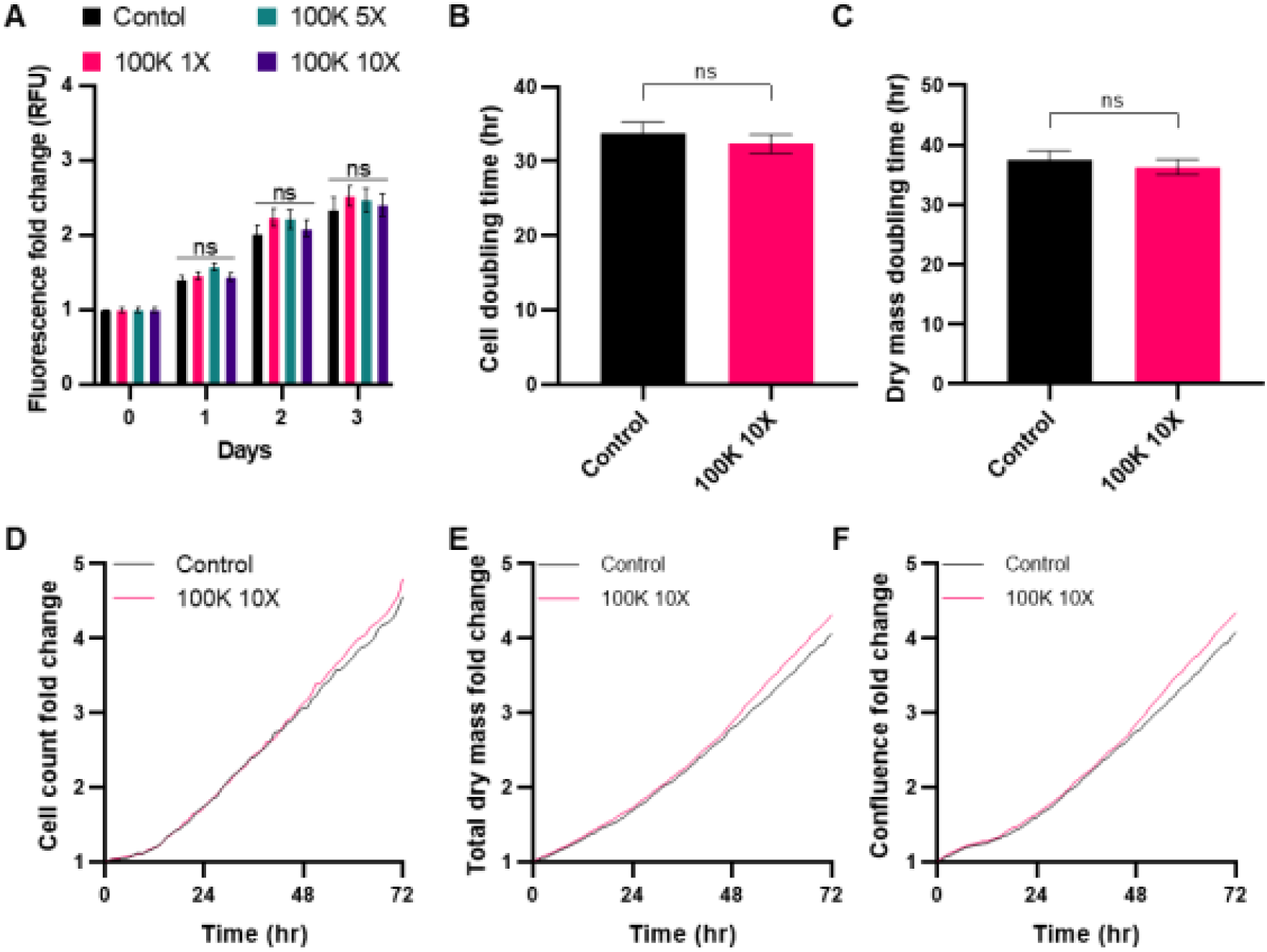
Effect of Y202 EVs on Y201 proliferation. A) Y201 cells treated with 1X, 5X and 10X concentrations of Y202 EVs and cell number determined over 72hrs (CyQuant assay). Livecyte image analysis was used to determine the effect of 10X Y202 EVs on Y201 B) cell doubling time, C) dry mass doubling time, D) cell counts, E) dry mass, and F) Y201 confluence. n=3, t-test or Two-Way ANOVA, ns = not significant.

### Y201 EVs promote the migration of Y202 cells

Scratch wound assays were used to test the effect of Y201 EVs on Y202 cell migration using LiveCyte image analysis. We demonstrated that Y201 EV exposure induced a significant increase in wound gap closure compared to untreated controls (Figure 8A, B). The track speed of cells at the leading edge of the scratch wound was also significantly increased following EV exposure compared to controls (Figure 8C). The time needed to cover 50% of the gap area was significantly decreased by 9hrs and the collective migration of the cells (a measure of how fast the cells on the edge move, assuming that cells in both edges move at the same rate) was increased by approximately 2µm/hr with Y201 EV treatment compared to controls (Figure 8D, E). Single cell analysis employing the MTrackJ and chemotaxis plugins in ImageJ were also performed. These analyses demonstrated that the track length, velocity and forward migration index of treated cells were significantly increased with Y201 EVs compared to the untreated group (Figure 8F-H). Y202 EVs had no effect on Y201 migration (Supplementary Figure S6).

**Figure 8.**
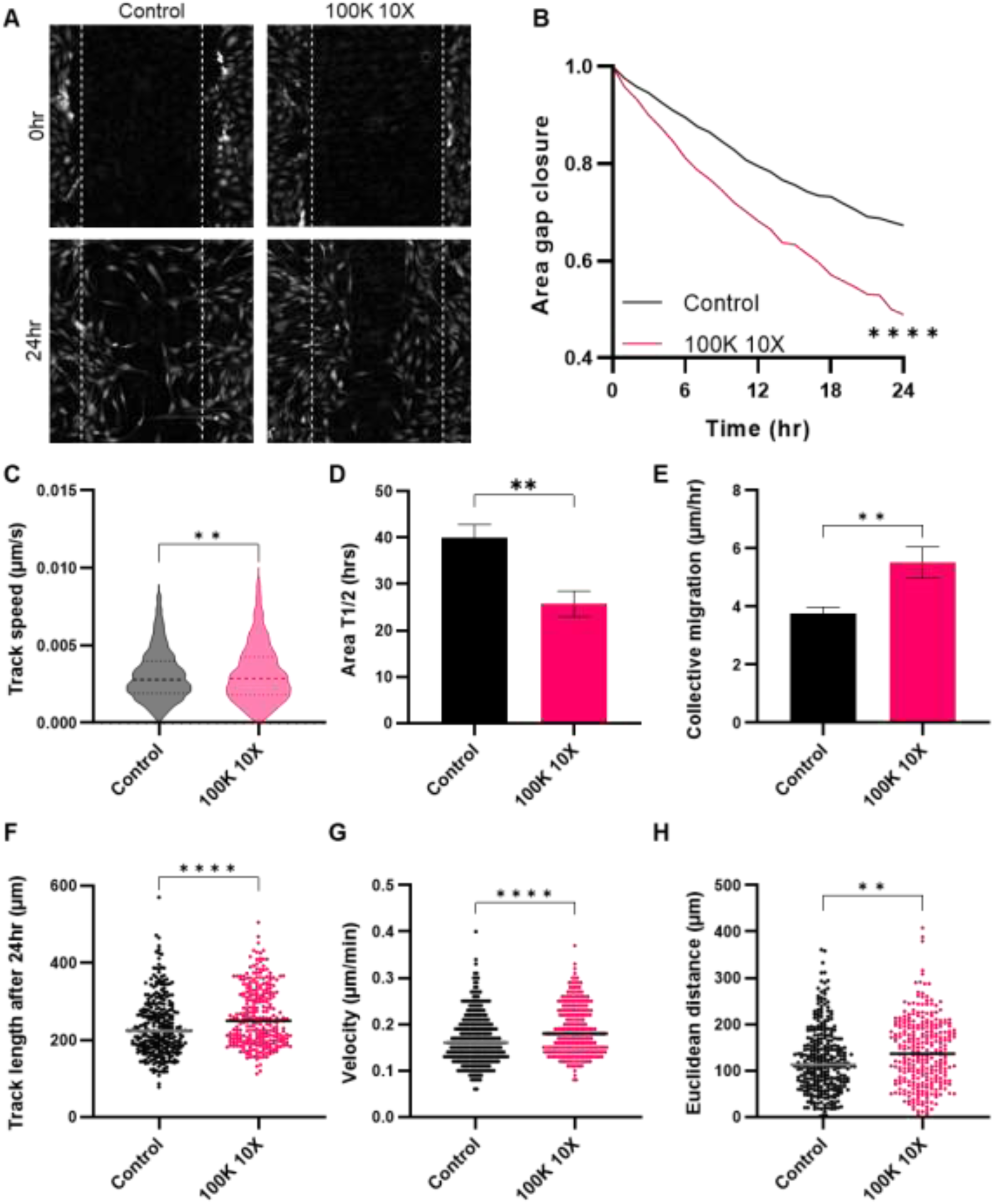
Effect of Y201 EVs on migration of Y202 cells. The scratch wound assay was used to monitor Y202 migration with and without exposure to Y201 EVs (100K 10X) using Livecyte image analysis. A) Micrographs of untreated control (left) and Y201 EV-treated (right) scratch-wounds at 0hrs (top) and 24hrs (bottom). Effect of Y201 EVs on B) gap area, C) Y202 track speed, D) area T1/2, E) Y202 collective migration, F) Y202 track length, G) cell velocity and H) Euclidean distance compared to untreated controls. n=3, t-test or Two-Way ANOVA *p<0.05, **p<0.01, ***p<0.001, ****p<0.0001.

### Effect of Y201 EVs on articular chondrocyte proliferation

The proliferative effect of Y201 EVs was also examined using primary articular chondrocytes (ACs) isolated from three samples from osteoarthritis (OA) patients (K268AC P3, K269AC P4, K272AC P3). The ACs were treated with 10X and 20X concentrations of Y201 EVs and proliferation was monitored for 72hrs using ptychographic quantitative phase imaging. ACs treated with Y201 100K EVs resulted in a significant increase in total cell counts, total dry mass and confluency over 72hrs in a dose-dependent manner (Figure 9A-C). The effect was particularly prominent during the final 24hrs of the experiment for the 20X treatment and the final 12hrs for the 10X treatment. AC doubling time was reduced by approximately 33hrs and by 43hr for 10X and 20X treatments respectively compared to the controls. AC dry mass was reduced by 107hrs and 157hr for 10X and 20X treatments respectively compared to the controls (Figure 9D-E). Y202 EVs had no significant effect on any measure of AC proliferation (Supplementary Figure S7).

**Figure 9:**
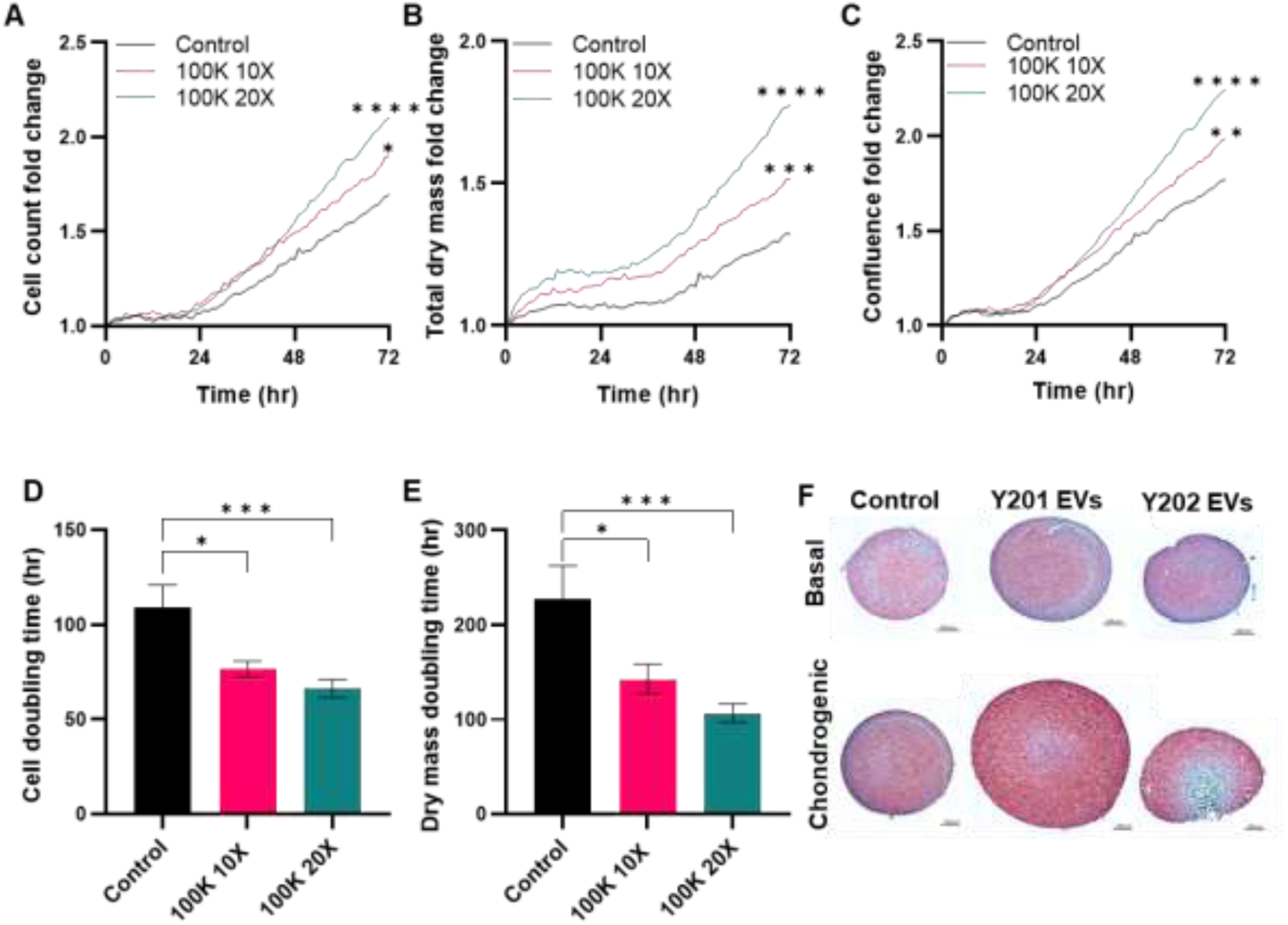
Effect of Y201 EVs on proliferation of primary articular chondrocytes and chondrogenesis. Primary articular chondrocytes were exposed to Y201 100K EVs at 10X and 20X concentrations. Cell proliferation was monitored by LiveCyte imaging over 72hrs determining A) cells counts B) total dry mass, C) confluence, D) cell doubling time and E) dry mass doubling time. n=3 Two Way-ANOVA followed by Tukey multiple comparison *p<0.05, **p<0.01, ***p<0.001, ****p<0.0001. F) Primary donor MSCs as micromass pellets in basal or chondrogenic differentiation conditions for 7 days with or without exposure to Y201 EVs and Y202 EVs. Micromasses were fixed, sectioned and stained with Safranin O. Red staining indicates chondrogenic differentiation. Image shows representative staining in one primary donor.

Considering these observations, we evaluated the chondrogenic effect of the Y201 and Y202 EVs. Micromass pellets were formed using primary MSCs, treated with Y201 and Y202 EVs and cultured in basal or chondrogenic differentiation medium for 7 days. Following histological staining with Safranin O, which identifies cartilage proteoglycans, we demonstrated that Y201 EV-treated pellets were significantly larger with more intense Safranin O staining compared to Y202 EVs and untreated controls (Figure 9F).

### RGD-integrin mediated uptake and function of Y201 EVs

Our findings have shown subtype-specific biological effects of EVs derived from different clonal sources of MSCs, which may be linked to the intra- and extra-vesicular protein composition and uptake mechanisms by target cells. Considering our evidence for an enriched ECM-based corona in Y201 EVs, particularly the abundance of RGD-containing proteins (for example, FN1 and MFG-E8), we hypothesised that these EV subtypes were preferentially taken up by integrin-mediated endocytosis. Y202 cells were used as model target cells and treated with GRGDSP integrin blocking peptides or GRADSP peptide controls for 6hrs followed by exposure to CFSE stained Y201 EVs. Using flow cytometry, we demonstrated that Y201 EV uptake was significantly inhibited by RGD blockade compared to controls (Figure 10A, B). Focal adhesion kinase (FAK) acts downstream of RGD integrins to mediate outside-in signalling through phosphorylation of ERK1/2 (Legate, Wickström and Fässler, 2009). We demonstrated by western blot analysis that Y201 EVs stimulated phosphorylation of ERK1/2 after 30 mins of exposure in Y202 cells, and that this effect was mediated by FAK (Figure 10C-F).

**Figure 10:**
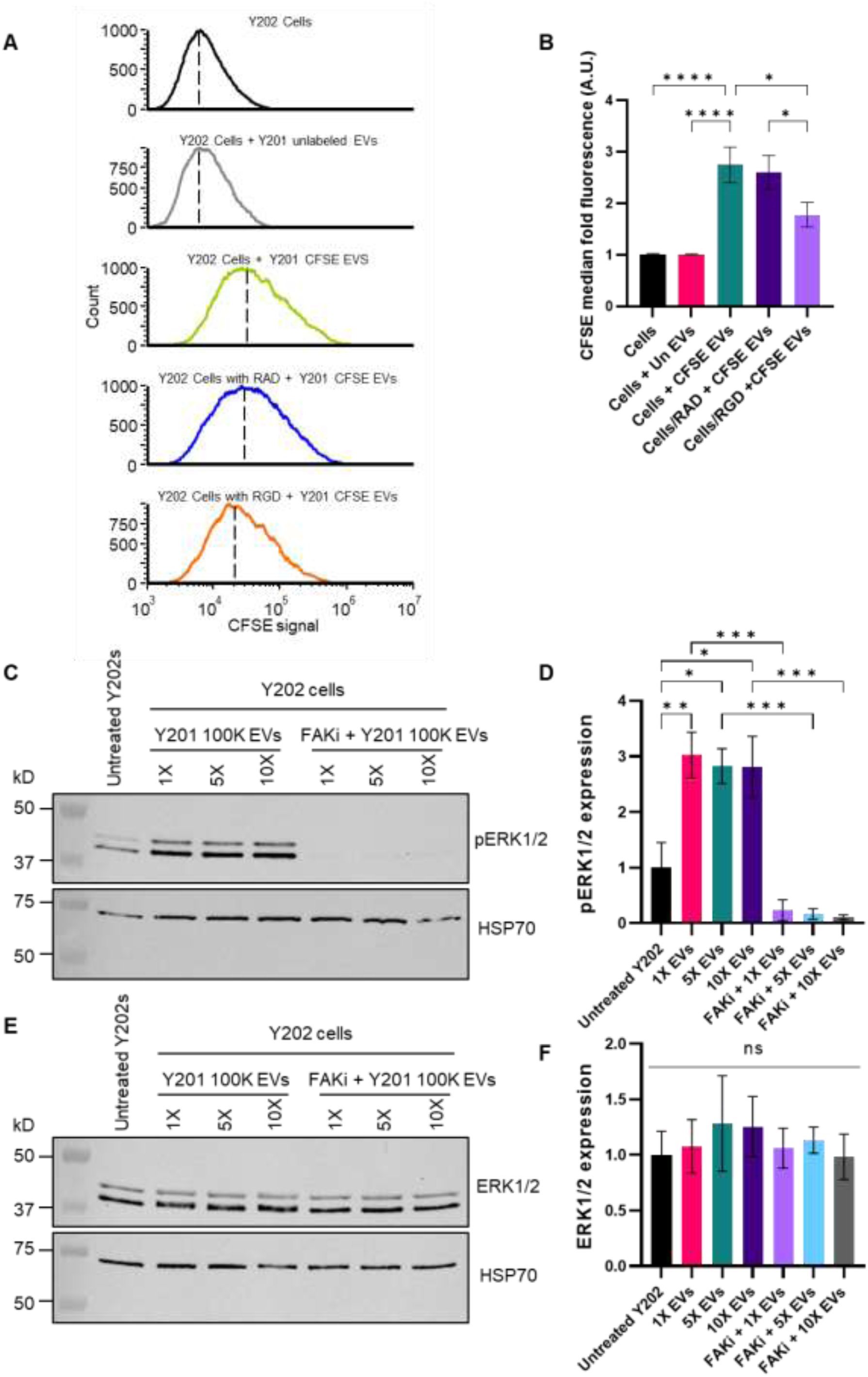
Mechanistic analysis of Y201 EVs. The uptake of CFSE-labelled Y201 EVs cells, with and without RGD blocking peptides (GRGDSP) or RAD control peptides (GRADSP), and effect on phosphorylation of ERK1/2 (pERK1/2) was determined in Y202 cells. A) Histograms showing CFSE signal in Y202 cells after exposure to CFSE-labelled Y201 EVs for 4hr. Green histogram shows EV uptake in untreated control Y202 cells; blue histogram shows EV uptake in the presence of RAD peptides; orange histogram shows EV uptake in the presence of RGD peptides. B) Fold change quantification of the CFSE median fluorescence of the Y202 cells, n=6. C-F) Western blot analysis of the effect of Y201 EVs on phosphorylation of ERK1/2 (pERK1/2) (C&D) and total ERK1/2 (E&F) in Y202 cells. Y202 cells were treated with Y201 EVs for 30mins at various concentrations (1X, 5X and 10X) in the absence and presence of a focal adhesion kinase inhibitor (FAKi) and western blotting (C&E) with quantification by densitometry (D&F) was used to determine pERK1/2 and total ERK1/2 expression compared to HSP70 loading controls, n=3. Error bars = SEM, One-Way ANOVA with Bonferroni corrections, *p<0.05, **p<0.01, ***p<0.001, ****p<0.0001.

To confirm the role of RGD integrins in Y201 EV bioactivity, primary ACs were treated with GRGDSP (and the GRADSP control peptide), and the effect of Y201 EVs on their proliferation rate was monitored for 72hrs. We confirmed that Y201 EVs increased AC proliferation compared to untreated controls and that this effect was blocked by exposure to GRGDSP but not GRADSP peptides (Figure 11).

**Figure 11:**
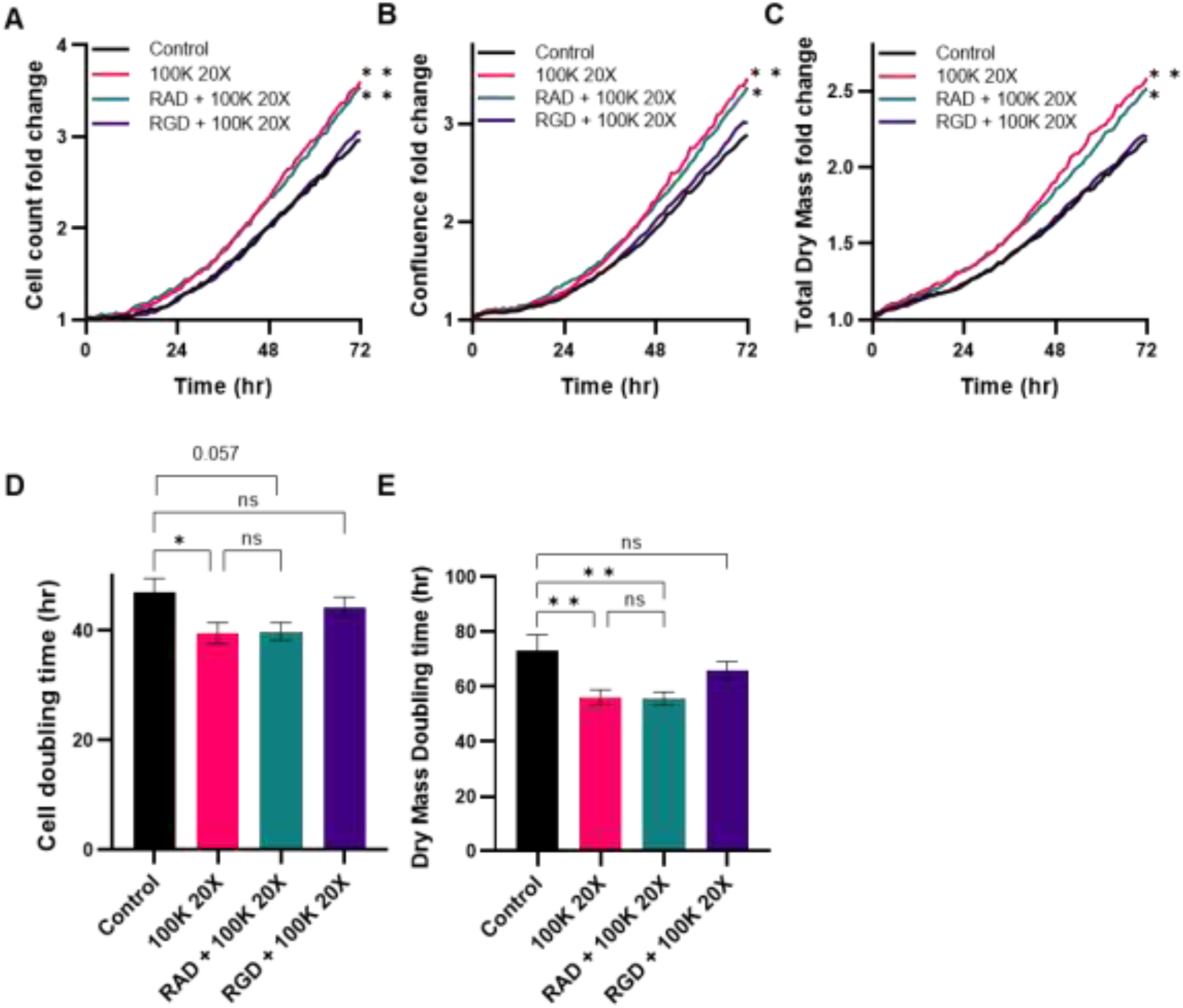
Effect of Y201 EVs on proliferation of primary articular chondrocytes, with and without RGD blocking peptides (GRGDSP) or RAD control peptides (GRADSP). A) cell counts, confluence, C) total dry mass, D) cell doubling time and E) dry mass doubling time of chondrocytes. n=3, Error bars= SEM, Two-Way or One-Way ANOVA, *p<0.05, **p<0.01, ns = not significant.

### Effect of Y201 EVs and Y202 EVs on T cell function

A frequently reported immune function of MSCs is to suppress activated T cell proliferation. Here, T cell proliferation was assessed by determining gradual division (proliferative index) and population doublings (proliferative cycles) over 5 days of co-culture with or without Y201 EVs and Y202 EVs. Compared to T cells alone, both EV subsets significantly reduced proliferative index scores and Y202 EVs reduced proliferative cycles (Figure 12A, B). Next, we determined the influence of Y201 EVs and Y202 EVs on the polarisation of naïve T cells into effector lineages with immunosuppressive/anti-inflammatory function. Y201 EVs, but not Y202 EVs, caused a significant increase in the development of anti-inflammatory Th2 cells, as indicated by intracellular IL4 staining (Figure 12B). There were no significant effects on Th1 Th17 and Treg cell numbers, as indicated by IFN-γ, IL17a and CD25/FOXP3 expression respectively (Figure 12C).

**Figure 12.**
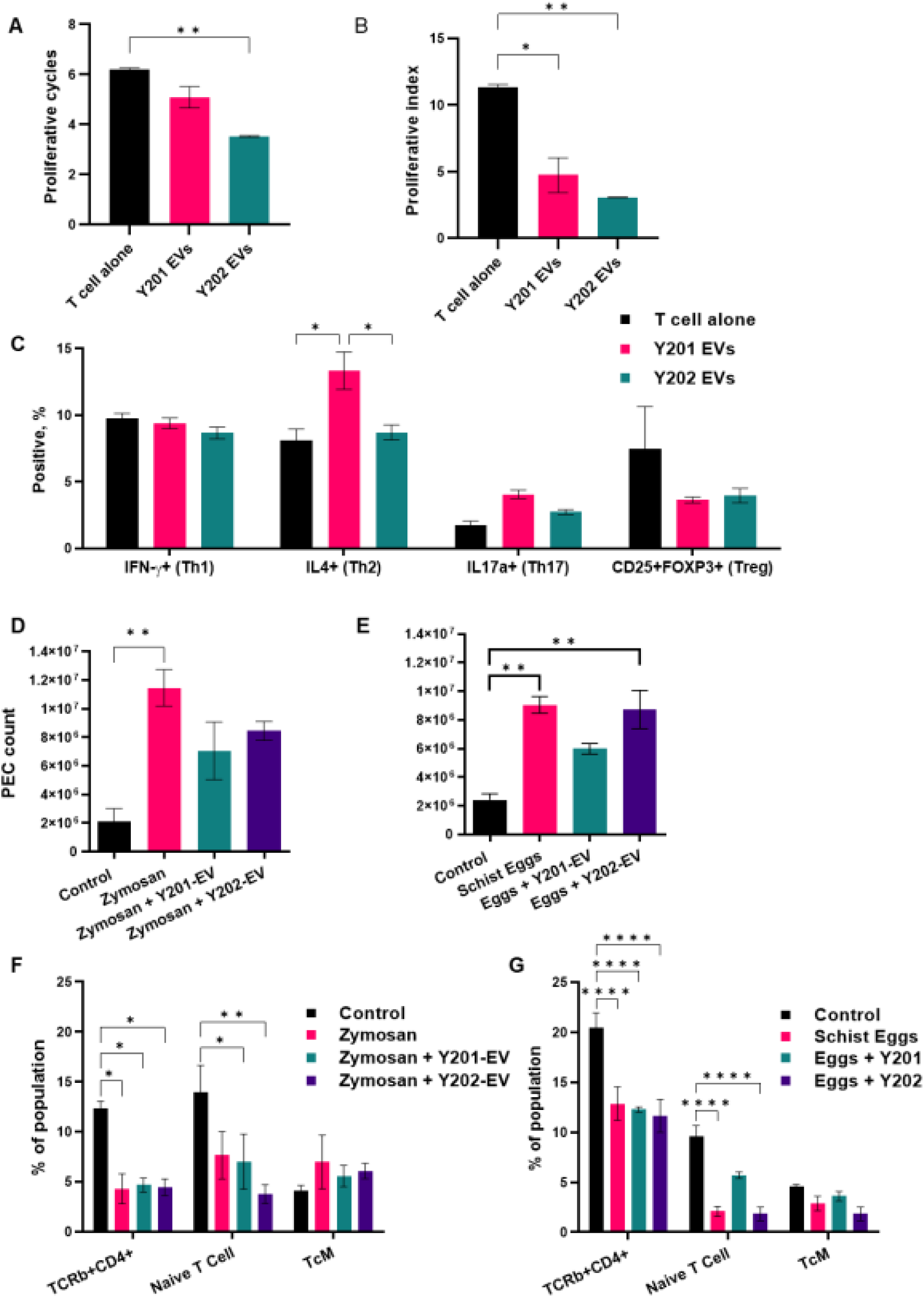
Immunomodulatory capacity of EVs isolated from Y201 and Y202 MSC subtypes. In vitro treatment of activated CD4+ T cells with Y201 and Y202 EVs (n=2, mean events >18,000 counted) A) Proliferative cycles. B) Proliferative index C) Polarisation of activated T cells in the absence and presence of Y201 EVs and Y202 EVs (n=2, mean events >32,000 counted). D/E) Total peritoneal exudate cell (PEC) counts following zymosan (D) or schistosome egg (E) induced inflammation in the absence and presence of Y201 EVs and Y202 EVs. F/G) Examination of TCR+CD4+, naïve and central memory T cells in zymosan or schistosome egg induced inflammation in the absence and presence of Y201 EVs and Y202 EVs (n=3). One-Way ANOVA with Bonferroni post hoc testing,*p<0.05, **p<0.01, ***p<0.001.

### *In vivo* assessment of immunomodulatory capacity of Y201 and Y202 EVs in a murine peritonitis model

Immune regulation was evaluated by Y201 and Y202 MSCs in zymosan- and schistosome egg-induced peritonitis model of acute inflammation that promotes the recruitment of monocytes and neutrophils to the peritoneal cavity. Following treatment with irritant, peritoneal exudate cells (PEC) were collected by lavage and analysis performed on the cell content. A gating strategy was devised for flow cytometric analysis of multiple PEC cell types focusing on haematopoietic, myeloid and lymphoid cells including monocytes, macrophages and T cells. In contrast to untreated zymosan irritant (11.5 ± 1.3 x 10^6^ cells), treatment with EVs from either Y201 (7.1 ± 2.0 x 10^6^ cells) or Y202 (8.5 ± 0.7 x 10^6^ cells) MSC lines did not significantly increase the number of PEC cells in comparison to PBS controls without inflammation (2.1 ± 1.6 x 10^6^ cells) (Figure 12D). When schistosome egg irritant was applied, both untreated irritant (9.1 ± 0.6 x 10^6^ cells) and Y202 EVs treatment (8.7 ± 1.3 x 10^6^ cells) showed significant recruitment of immune cells to the peritoneal cavity over PBS controls (2.4 ± 0.4 x 10^6^ cells) whilst Y201 EVs (6.0 ± 0.4 x 10^6^ cells) suppressed increases observed in PEC counts (Figure 12E). It was notable that using the schistosome egg-induced peritonitis model, Y201 EV were able to maintain naive T cell numbers at levels significantly higher than egg-treatment only or Y202 EVs (Figure 12 F, G). Similarly, Y201 EV treatment dampened the effects of the irritant on myeloid cell recruitment to the region of inflammation, typically seen in acute *Schistosoma mansoni* infection. In particular, data indicated suppression of macrophage and neutrophil recruitment to the peritoneal cavity in response to schistosome egg-induced peritonitis (Supplementary Figure S8).

### *In vivo* assessment of potential therapeutic efficacy of Y201 and Y202 EVs in a murine arthritis model

Finally, we tested the bioactivities of Y201 and Y202 EVs in a disease-relevant *in vivo* model of adjuvant-induced arthritis model following intra-articular EV injection. Following histological examination and blind scoring, we demonstrated that EVs derived from Y201 MSCs induced a significant decrease in all measures of disease activity compared to vehicle controls, including joint swelling, synovial infiltrate, joint exudate, synovial hyperplasia and overall arthritis index (Figure 13 A-E). Y202 EVs significantly reduced joint swelling compared to controls but otherwise did not affect any other disease score measures. Representative haematoxylin and eosin-stained sections provide evidence of changes in synovial infiltrate and hyperplasia in control, Y201 EV and Y202 EV-treated samples (Figure 13 F-K).

**Figure 13.**
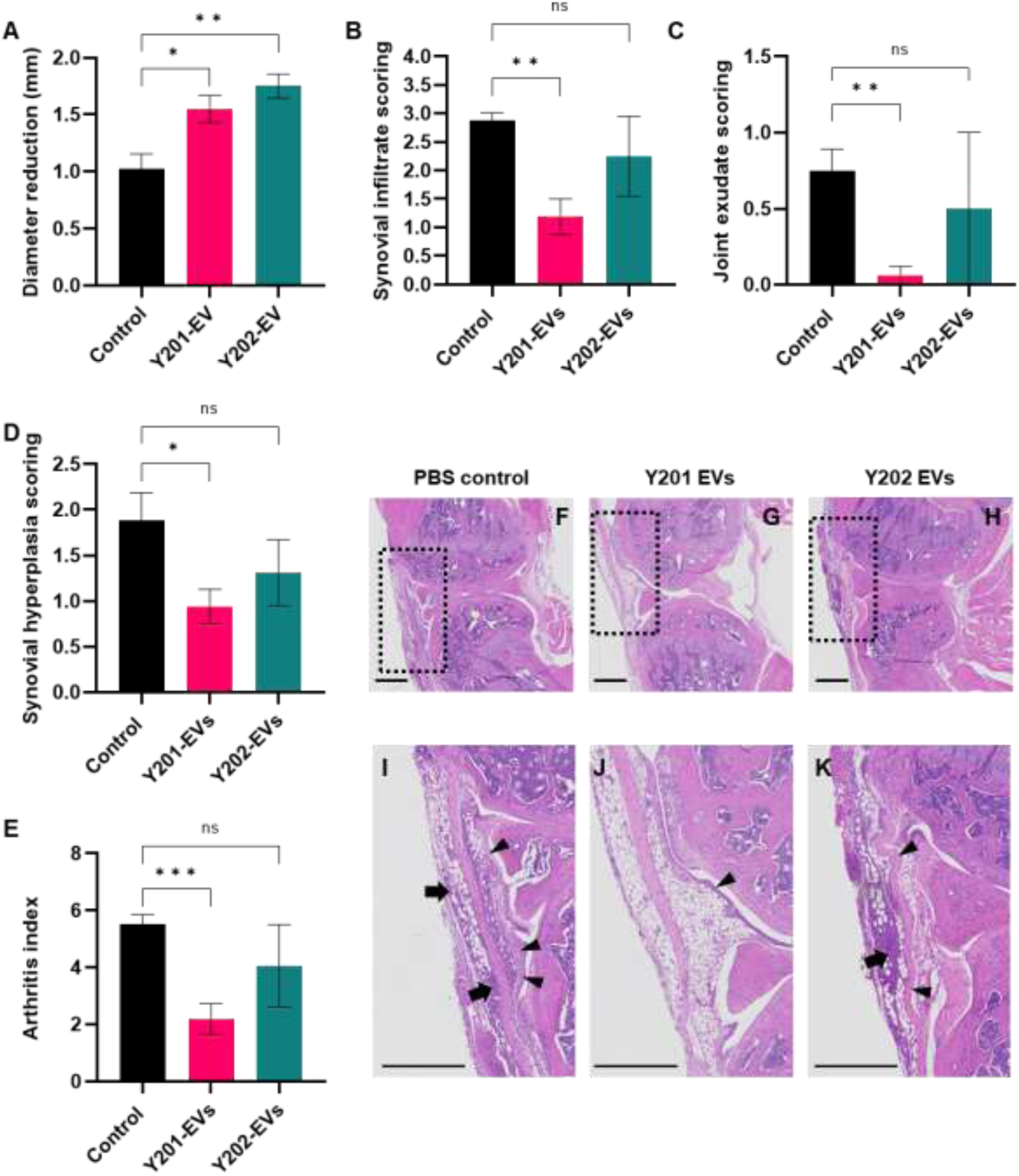
Effect of Y201 and Y202 EVs on disease activity in an in vivo adjuvant-induced arthritis model. A) Joint size reduction from peak swelling (mm), B) synovial infiltrate score, C) joint exudate score, D) synovial hyperplasia score, E) arthritis index. Haematoxylin and eosin-stained sections of F and I) PBS control, G and J) Y201 EV-treated mouse knee joints, and H and K) Y202 EV-treated mouse knee joints. Boxed areas in F, G and H are enlarged in I, J and K respectively showing examples of synovial infiltrate (cellular infiltration into synovium; arrows) and synovial hyperplasia (thickened synovial lining; arrowheads). Scale bars = 500mm. Mean values shown ± SD (n=4) Brown-Forsythe ANOVA test, *p<0.05, **p<0.01, ***p<0.001, ns = not significant.

## Discussion

There is a growing interest in the therapeutic application of EVs derived from different cell sources and MSCs in particular. It is clear that EV composition and bioactivity is determined to a large extent by the biology of the parent cell (Hurwitz *et al*., 2016; Sork *et al*., 2018; Hagey *et al*., 2023) and MSC tissue source (Cai *et al*., 2020; Almeria *et al*., 2022). Here we have demonstrated that stromal cell lines, clonally isolated from the same mixed donor population of MSCs, have distinct features and functionalities. These findings are important as almost all basic research, preclinical and clinical studies employ EVs from heterogeneous MSC pools, which we can now confirm will be prone to significant within-population variability. Moreover, the analysis of EVs derived from two related MSC subtypes with different bioactivities also assisted the identification of key functional EV features, which has informed MSC EV mechanism of action.

It was notable that the repertoire of lipids, miRNAs and proteins presented by Y201- and Y202-derived EVs was remarkably similar and may contribute to defining a common stromal EV biomarker signature. More in-depth analyses demonstrated differences in the relative abundance of these EV components, particularly within the EV proteome, where Y201 EVs were found to be significantly enriched compared to Y202 EVs. Numerous studies have analysed the MSC-EV proteome with a diverse array of outcomes, comprehensively reviewed by Qiu and colleagues (Qiu *et al*., 2019). A detailed interrogation of published reports has described an MSC-EV-specific signature, by comparing 10 MSC-EV datasets with 12 non-MSC-EV datasets (van Balkom *et al*., 2019). The signature includes a number of proteins we identified, including collagen I and VI, filamins, actin and vimentin, but noticeably absent were FN1 and MFG-E8, which are the two most abundant Y201-EV proteins. Although the Y201-EVome is varied and complex, the enrichment of RGD-containing FN1 and MFG-E8 may be a defining feature that directly contributes to their bioactivity. Bioinformatic interrogation of our own datasets demonstrated that Y201 EV proteins predominantly contribute to matrix organisation, cell-cell and cell-matrix interactions as well as immune regulation. We hypothesise that a significant proportion of the identified proteins contribute to an EV corona, presented on the exofacial surface by integrins and other adhesion molecules, though further experimental validation is required. There is a growing understanding of the influential contribution the corona makes to EV function (Tóth *et al*., 2021; Buzas, 2022; Hallal *et al*., 2022; Wolf *et al*., 2022). Recent work has shown that EV-delivered cargoes were only detectable in recipient cells at high EV doses and that the overriding biological effects were mediated by EV surface proteins at lower EV doses that more closely represented physiological conditions (Hagey *et al*., 2023).

When comparing Y201 EVs with Y202 EVs, we attempted to consider physiologically-relevant effects by adopting a dosing strategy that accounted for EV yield per cell (Y201 MSCs produce significantly more EVs per cell than Y202 MSCs), though we recognise that improved attempts to define EV dose in this study and others are required. We were able to demonstrate that the effects of Y201 EVs were mediated via an RGD (integrin)-FAK-ERK1/2 axis, with increases in pERK1/2 detected within 30 minutes of exposure. Similar observations have been reported in other cell types (Purushothaman *et al*., 2016; Shtam *et al*., 2019; Fuentes *et al*., 2020) though the vast majority of studies focus on EV cargo delivery as a means of mediating biological effect. Here, we showed that Y201 EV internalisation, and therefore cargo delivery, took 2-4 hours. These findings demonstrate that target cell responses will be determined not only by biological make-up of EVs presented to the cell surface, but also the cell’s capacity for EV uptake.

The precise mechanisms by which Y201 EVs exert anti-inflammatory effects *in vitro* and *in vivo* are less clear, though MFG-E8 is an interesting candidate. MFG-E8 (also known as lactadherin) is an anti-inflammatory glycoprotein, enriched in the milk fat globule membrane (MFGM) where it contributes to its immune-boosting and antipathogenic activity to protect newborns (Peterson *et al*., 2013). Interestingly, lactosylceramide, which was significantly enriched in Y201 EV lipids, is also abundant in the MFGM (Ma *et al*., 2020). MFGE8 has protective effects in rheumatoid arthritis and other inflammatory conditions (Aziz *et al*., 2011; Albus *et al*., 2016; Yi, 2016). MFG-E8 can also restore chondrocyte function and reduce inflammatory bone loss in osteoarthritis (Abe *et al*., 2014; Lu *et al*., 2021). It is tempting to speculate that Y201 EVs mimic an apoptosis-induced anti-inflammatory response whereby Y201 EVs, like apoptotic cells, present MFG-E8 to immune cells to promote cell clearance and suppress inflammation (Yi, 2016). Intriguingly, our findings may explain why the immunosuppressive effects of infused MSCs appear to be mediated by MSC apoptosis (Galleu *et al*., 2017). Further work will be required to define these mechanisms with greater authority, but current evidence suggests that EVs derived from a population of Y201-type MSCs would have high therapeutic value in the treatment of inflammatory disorders associated with tissue loss.

## Acknowledgments

This work was funded by the Biotechnology and Biological Sciences Research Council United Kingdom, Doctoral Training Partnership grant (BB/M011151/1) and the Tissue Engineering and Regenerative Therapies Centre Versus Arthritis (21156). Instrumentation used is part of the York Centre of Excellence in Mass Spectrometry which was created thanks to a major capital investment through Science City York, supported by Yorkshire Forward with funds from the Northern Way Initiative, and subsequent support from engineering and physical sciences research council Grants (EP/K039660/1; EP/M028127/1). We thank Adam Dowle of the York Centre of Excellence in Mass Spectrometry for assistance with the LC-MS/MS.

