## Supplementary Materials for "Extracellular vesicle bioactivity and potential clinical utility is determined by mesenchymal stromal cell clonal subtype"

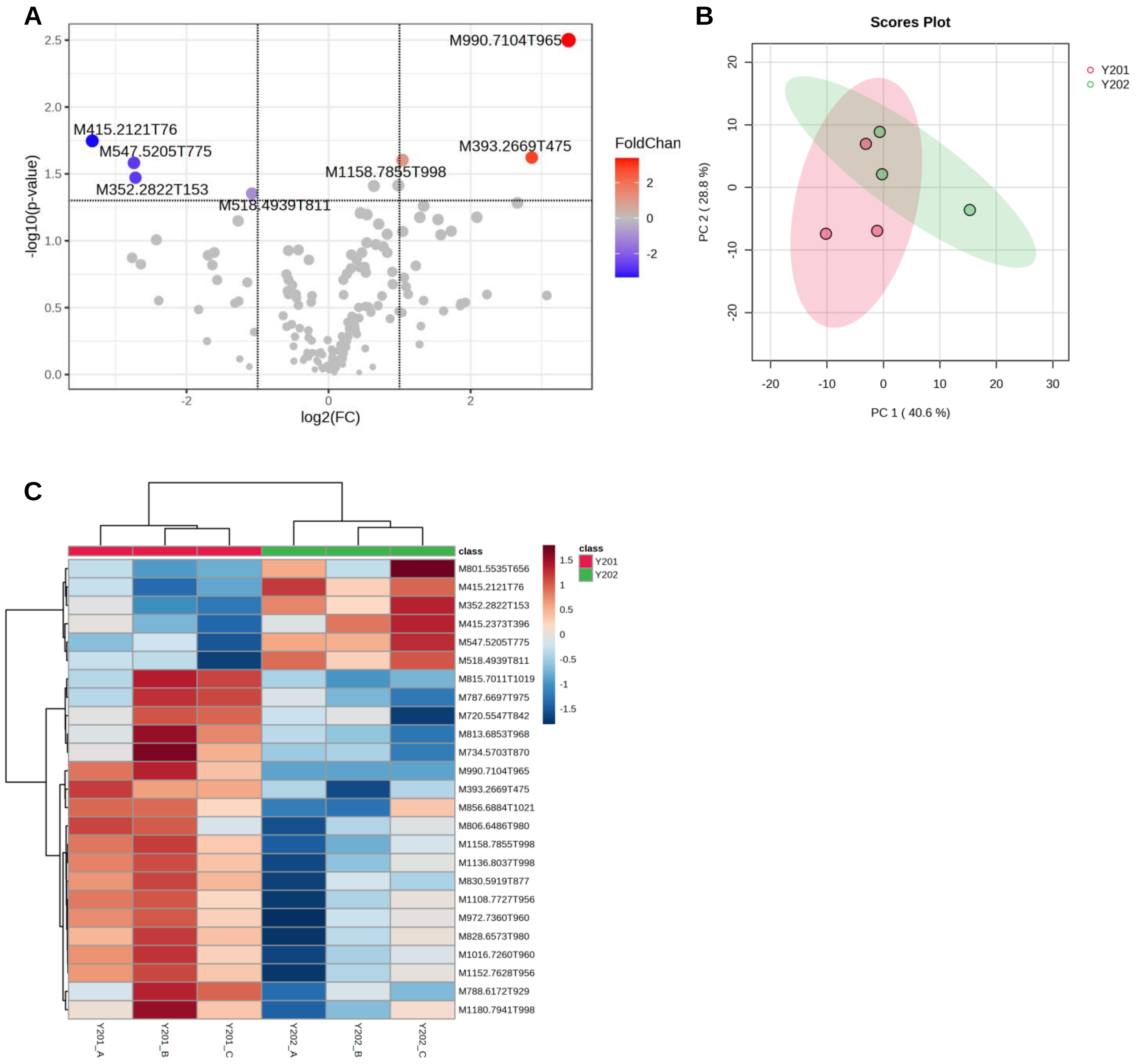


**Supplementary Figure S1. Analysis of Y201 and Y202 EV lipids.** A) Volcano plot of lipids identified in EV isolations from Y201 and Y202 clonal lines in a fold change format. 172 different lipids were identified and lipids with a fold change >2 were subjected to a t-test ($p<0.05$). The two upper quadrants contain lipids that were significantly different between Y201 and Y202 EVs. The blue dots indicate lipids that are downregulated in Y201 EVs and the red dots the upregulated lipids. The analysis was performed relative to Y201 EVs. B) Principal components analysis (PCA) scores plots of Y201 vs Y202 EVs, shaded areas are the 95% confidence regions of each group, C) Hierarchical clustering heatmap analysis illustrating the lipidomic profiles of triplicate samples of Y201 EVs (red, left) and Y202 EVs (green, right). Each bar represents a lipid feature at average intensity on a normalised scale from blue (low) to red (high). The dendrograms were constructed using the Euclidian distance method with the ward clustering algorithm.

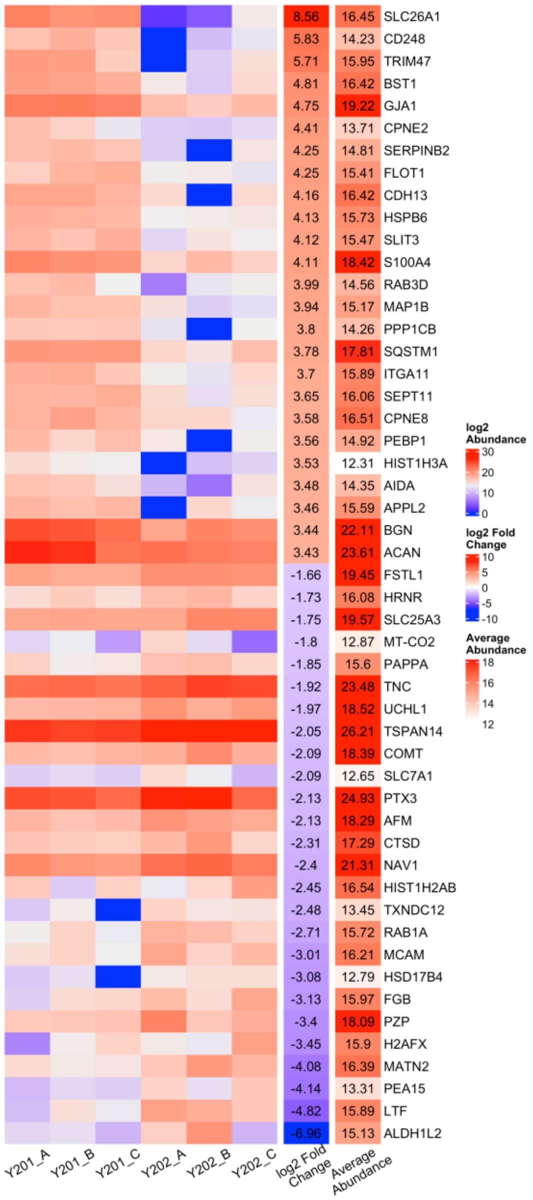


**Supplementary Figure S2.** Heatmap of protein abundance for the top and bottom 25 proteins with the highest fold change relative to Y201 EV (left); Log2 fold change (second right); Mean abundance across both Y201 and Y202 EV populations (far right)

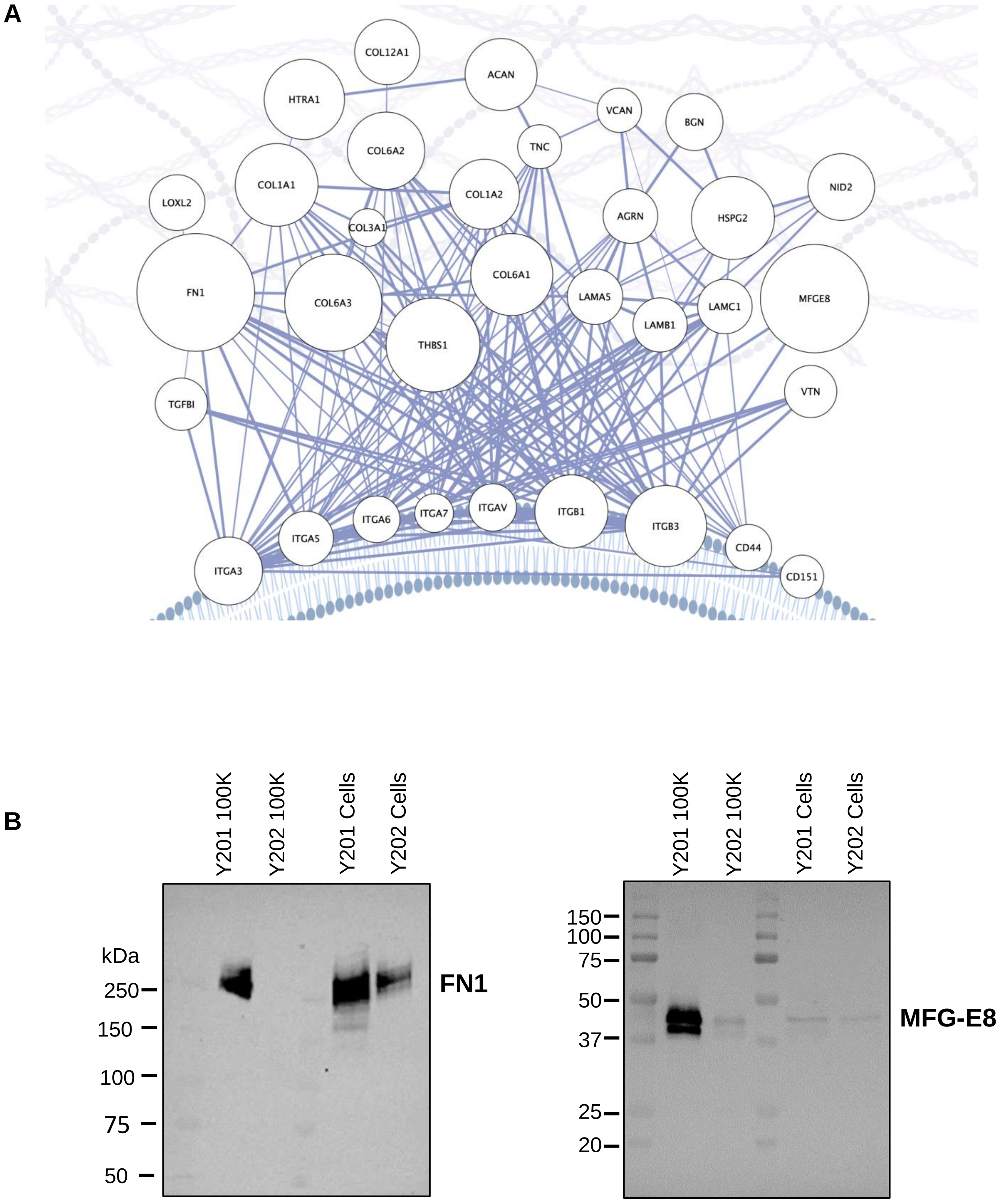


**Supplementary Figure S3**. A) STRING PPI network detailing relative abundance and strength of interactions between proteins in the Y201 core proteome predicted at the EV surface. Node size was mapped to the relative abundance of proteins; Edge width corresponds to combined score of the interaction. Network visualisation created using BioRender.com. B) Western blot analysis of Fibronectin-1 (FN1) and MFG-E8 in Y201 and Y202 EVs and whole cell lysates.

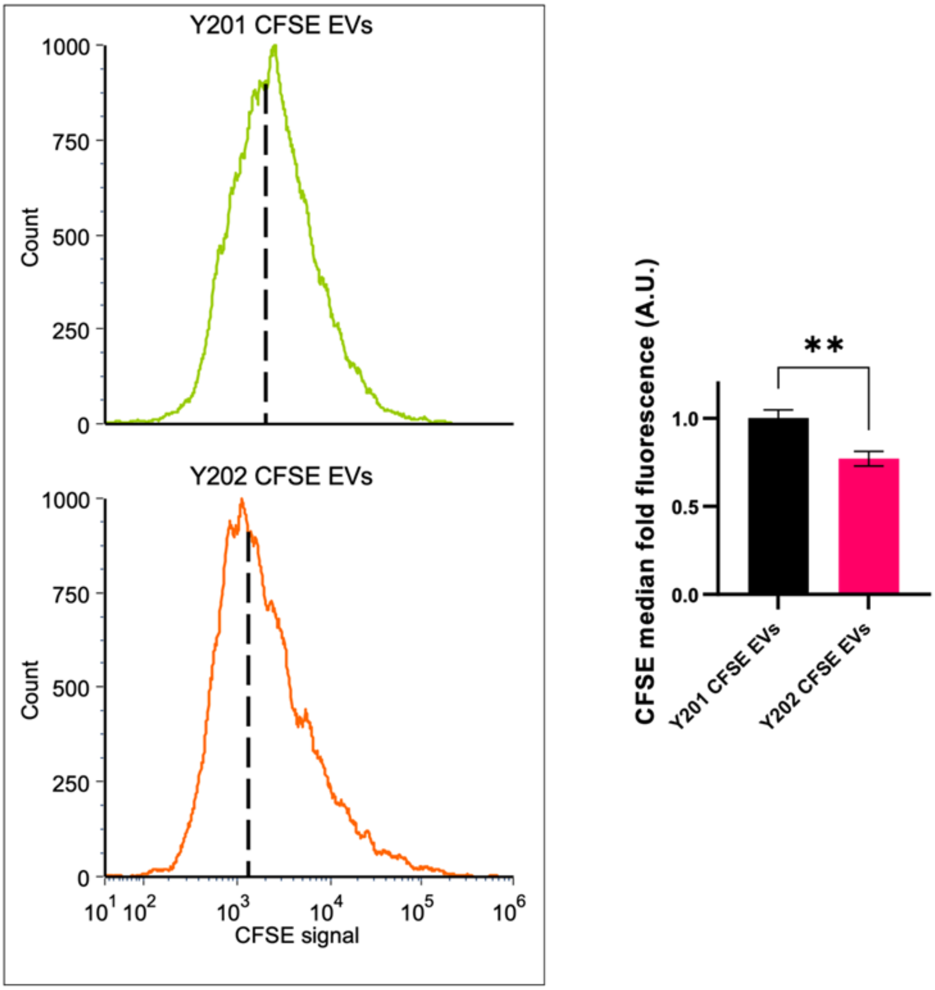


**Supplementary Figure S4. Flow cytometric analysis of CFSE-stained Y201 and Y202 EVs**. The fluorescence intensity of CFSE-EVs was quantified using the Cytoflex S system. Representative histograms (left) show the CFSE signal from both Y201 and Y202 CFSE EVs. The dashed lines represent the median fluorescence readings, quantified (right). n=4, t-test, **$p<0.01$

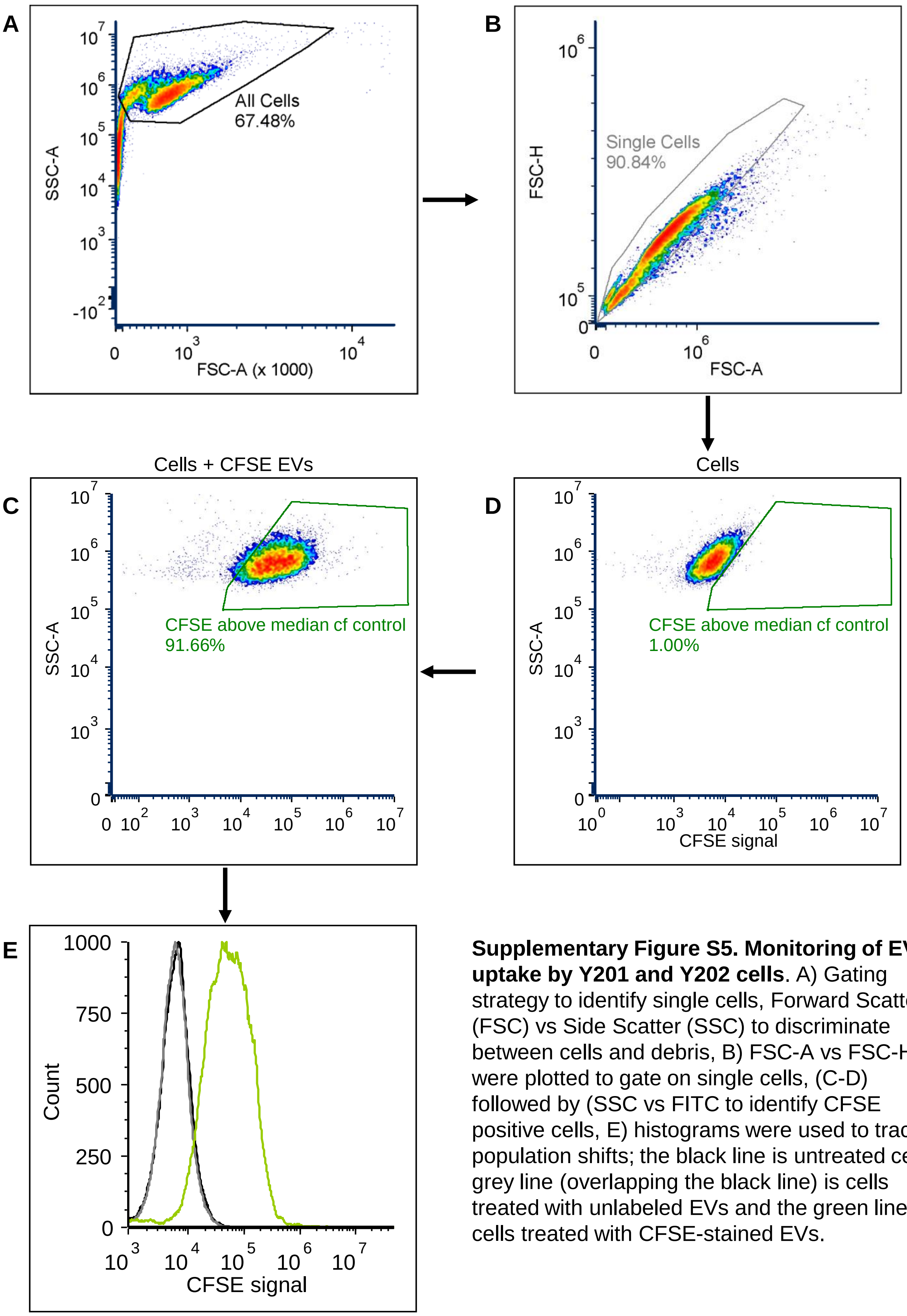


**Supplementary Figure S5. Monitoring of EV uptake by Y201 and Y202 cells**. A) Gating strategy to identify single cells, Forward Scatter (FSC) vs Side Scatter (SSC) to discriminate between cells and debris, B) FSC-A vs FSC-H were plotted to gate on single cells, (C-D) followed by (SSC vs FITC to identify CFSE positive cells, E) histograms were used to track population shifts; the black line is untreated cells, grey line (overlapping the black line) is cells treated with unlabeled EVs and the green line is cells treated with CFSE-stained EVs.

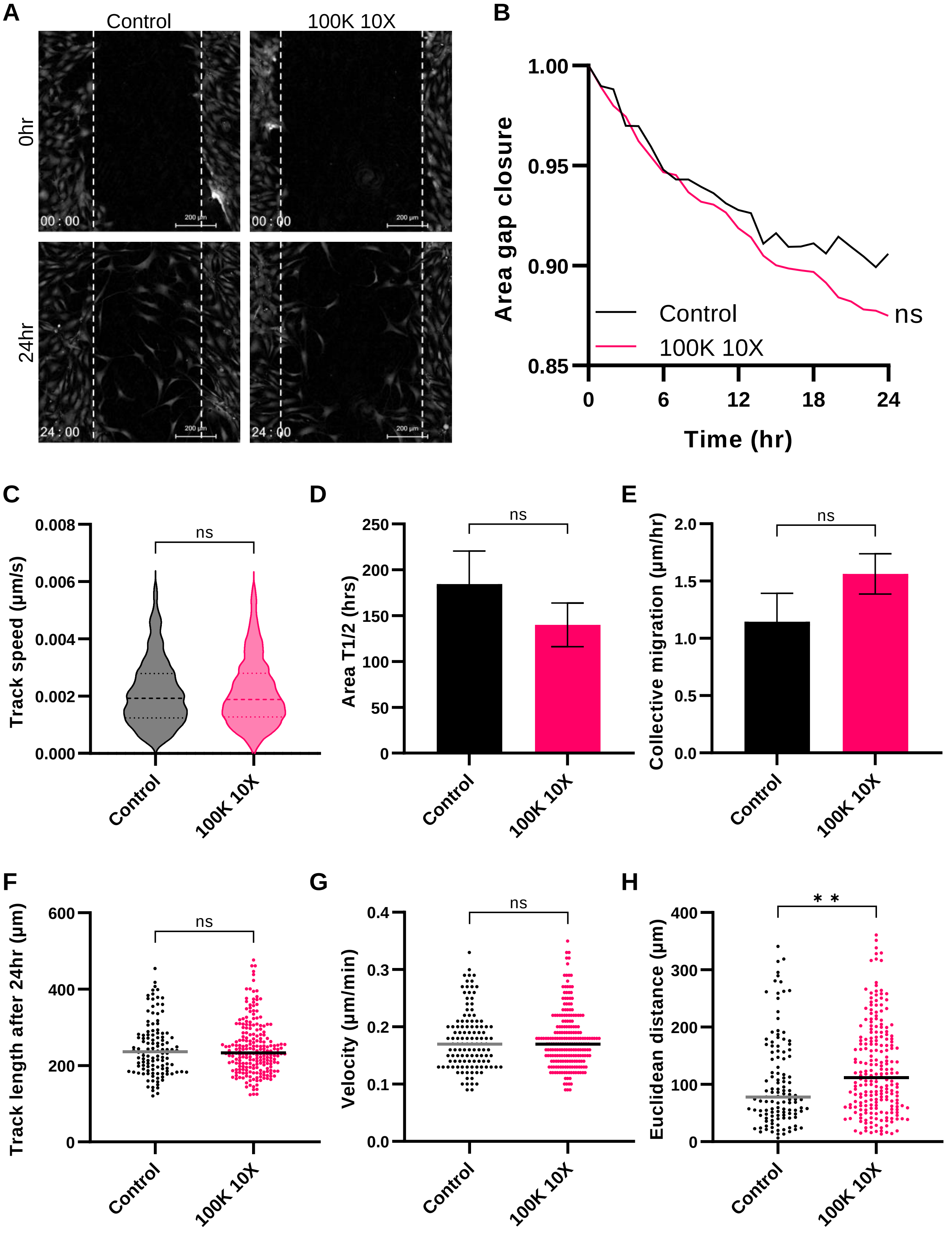


**Supplementary Figure S6. Effect of Y202 EVs on the migration of Y201 cells.** A scratch wound assay was used to monitor the migration of Y201 cells after they were treated with Y202 EVs (100K 10X) using LiveCyte image analysis. A) Micrographs of untreated control (left) and Y202 EV-treated (right) scratch at 0 hours (top) and 24 hours (bottom). Effect of Y202 EVs on B) gap area, C) Y201 track speed, D) area T1/2, E) Y201 collective migration, F) Y201 track length, **G)** cell velocity and **H)** Euclidean distance compared to untreated controls, n=2, t-test or Two-Way ANOVA, **p<0.01.

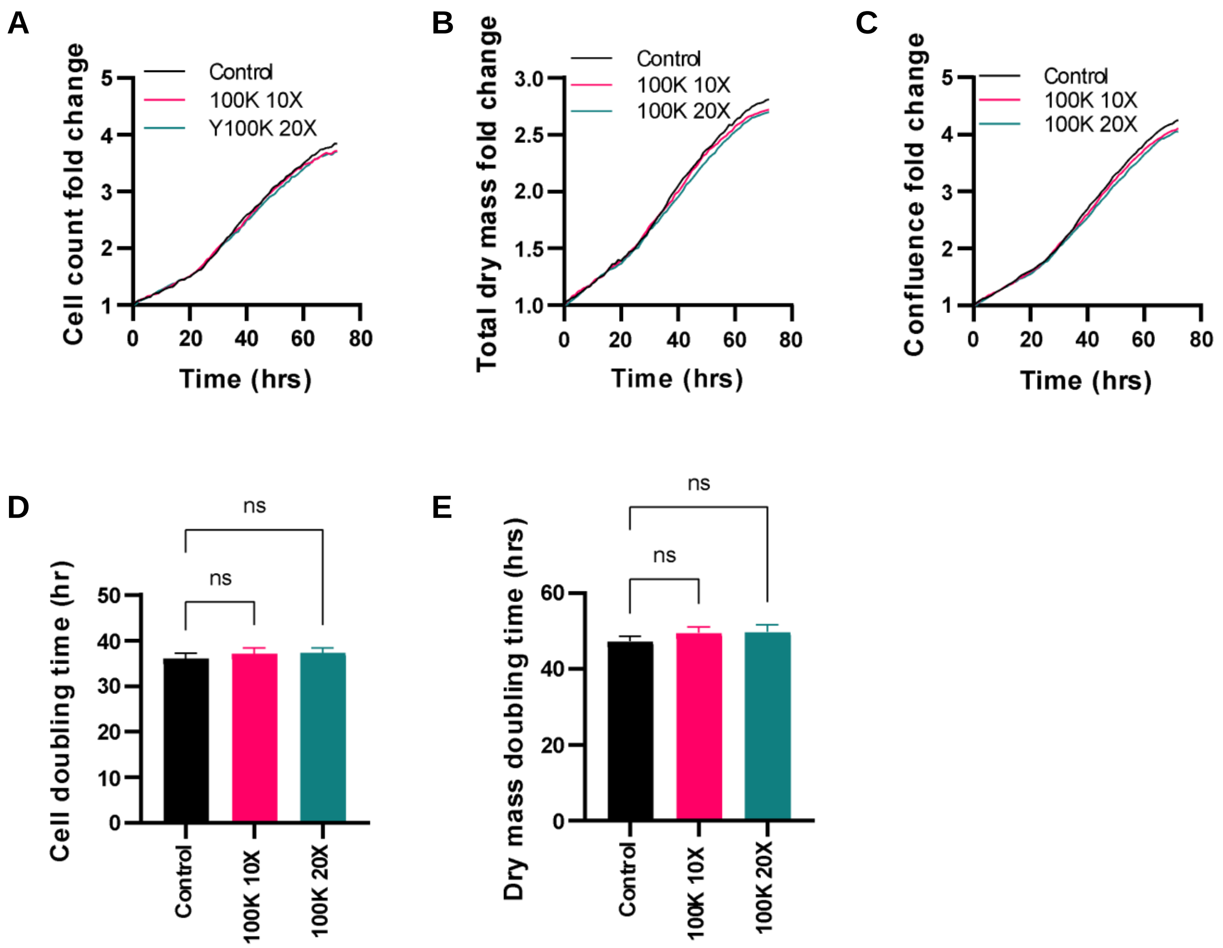


**Supplementary Figure S7: Effect of Y202 EVs on proliferation of primary articular chondrocytes.** Primary articular chondrocytes were exposed to Y202 100K EVs at 10X and 20X concentrations. Cell proliferation was monitored by LiveCyte imaging over 72hrs determining A) cells counts B) confluence, C) total dry mass, D) cell doubling time and E) dry mass doubling time. n=3, Two-Way ANOVA followed by Tukey multiple comparison, ns = not significant.

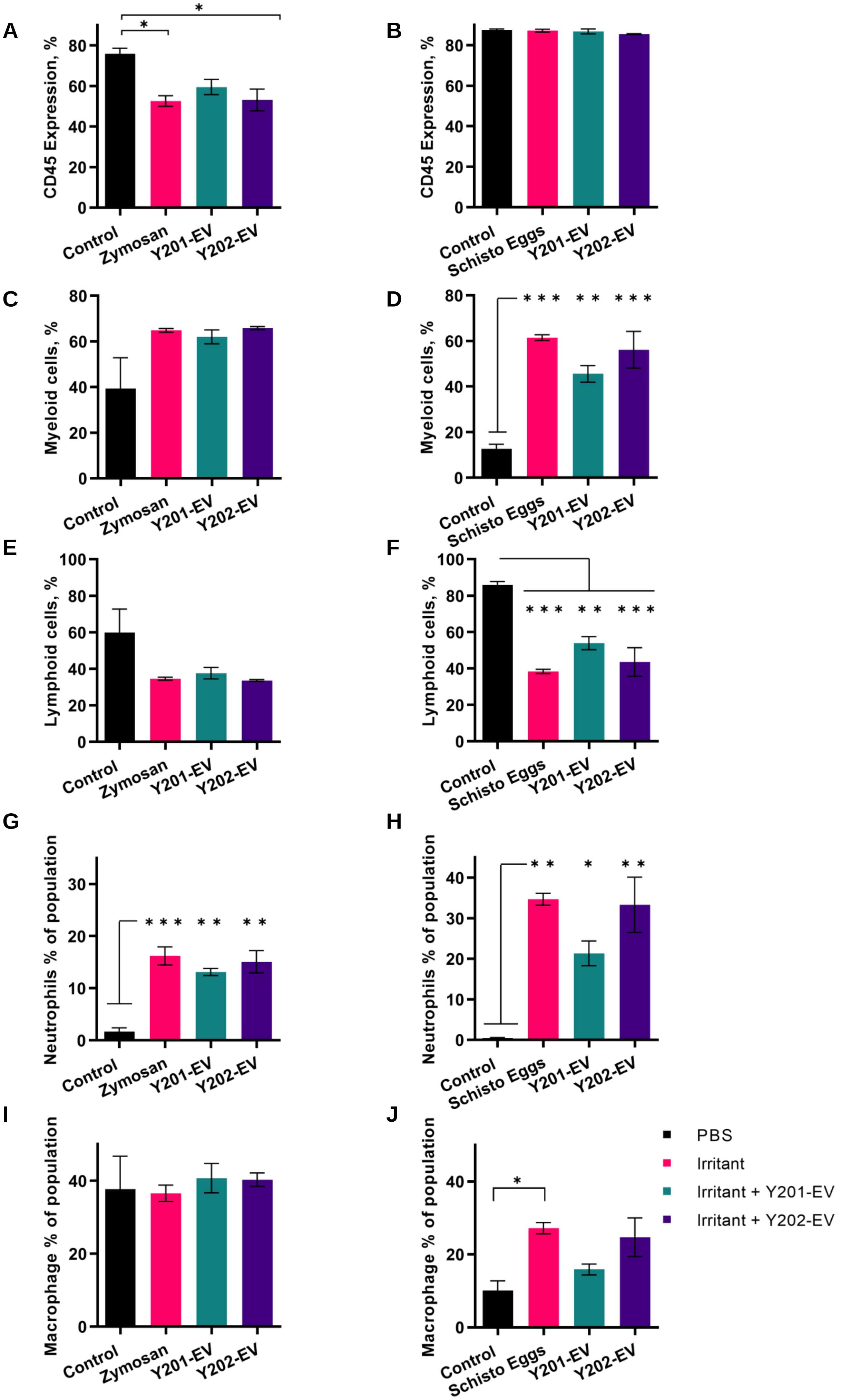

A
CD45 Expression, %
Control
Zymosan
Y201-EV
Y202-EV
B
CD45 Expression, %
Control
Schisto Eggs
Y201-EV
Y202-EV
C
Myeloid cells, %
D
Myeloid cells, %
E
Lymphoid cells, %
F
Lymphoid cells, %
G
Neutrophils % of population
H
Neutrophils % of population
I
Macrophage % of population
J
Macrophage % of population
PBS
Irritant
Irritant + Y201-EV
Irritant + Y202-EV

**Supplementary Figure S8. Y201- and Y202-derived EVs applied into an *in vivo* murine zymosan or schistosome egg induced peritonitis model of inflammation.** Flow cytometric analysis of peritoneal exudate fluid cells focused on haematopoietic, myeloid and lymphoid cells including neutrophils and macrophages. Quantified immune cell recruitment for Y201-derived EV treated peritonitis showed the presence of CD45+ (A/B); Myeloid (C/D) and Lymphoid (E/F) cells; Neutrophils (G/H); and Macrophages (I/J), n=3, One-Way ANOVA with Bonferroni post hoc testing,*p<0.05, **p<0.01, ***p<0.001

**Supplementary Table 1**. Relative fold changes for miRNAs identified as significantly different in Y201 and Y202 EVs.

| miRNA | Mean count Y201 (±SEM) | Mean count Y202 (±SEM) | Fold change Y201 vs Y202 |
|---|---|---|---|
| hsa-miR-376a-3p | 33.07 ± 6.23 | 8.49 ± 4.5 | 3.90 |
| hsa-miR-29a-3p | 91.92 ± 19.98 | 8.39 ± 4.82 | 10.96 |
| hsa-miR-221-3p | 65.9 ± 19.89 | 4.49 ± 1.04 | 14.68 |
| hsa-miR-145-5p | 56.1 ± 17.39 | 3 ± 0.8 | 18.70 |
| hsa-miR-630 | 85.3 ± 18.84 | 19.7 ± 11.95 | 4.33 |
| hsa-miR-21-5p | 138.21 ± 29.8 | 22.01 ± 10.38 | 6.28 |
| hsa-miR-100-5p | 90.94 ± 21.93 | 3.89 ± 0.6 | 23.38 |
| hsa-miR-320e | 30.11 ± 7.73 | 3 ± 0.8 | 10.04 |
| hsa-miR-125b-5p | 481.61 ± 94.95 | 145.49 ± 50.14 | 3.31 |
| hsa-miR-29b-3p | 67.84 ± 14.2 | 22.08 ± 10.63 | 3.07 |
| hsa-miR-612 | 27.29 ± 11.89 | 68.7 ± 11.1 | 0.40 |
| hsa-miR-6721-5p | 1.33 ± 0.28 | 24.47 ± 7.54 | 0.05 |

**Supplementary Table 2.** Summary of the most abundant lipid metabolites identified by lipidomic analysis in EVs derived from Y201 MSCs. Values were normalised by internal standard. Samples were analysed in the positive and negative ion modes of LC-MS/MS.

| Label | Annotation | polarity | Mean Y201 (±SD) |
|---|---|---|---|
| M760.5858T875 | PC(16:0_18:1) | pos | 19.11 ± 0.82 |
| M747.5657T801 | SM(d18:1/16:0) | neg | 18.47 ± 3.35 |
| M703.5755T801 | SM(d18:1/16:0) | pos | 16.01 ± 1.76 |
| M804.5762T874 | PC(16:0_18:1) | neg | 12.53 ±1.03 |
| M349.2402T395 | Prostaglandin | neg | 7.97 ±4.79 |
| M745.5502T739 | SM(d18:1/16:1) | neg | 5.53 ±1.12 |
| M788.6172T929 | PC(18:0_18:1) | pos | 5.09 + 0.96 |
| M786.6016T877 | PC(18:1_18:1) | pos | 4.83 ±0.60 |
| M701.5599T739 | SM(d18:1/16:1) | pos | 3.87 ± 0.82 |
| M830.5919T877 | PC(18:1_18:1) | neg | 3.79 + 0.40 |

**Supplementary Table 3.** Summary of the most abundant lipid metabolites identified by lipidomic analysis in EVs derived from Y202 MSCs. Values were normalised by internal standard. Samples were analysed in the positive and negative ion modes of LC-MS/MS.

| Label | Annotation | polarity | Mean Y202 (±SD) |
|---|---|---|---|
| M747.5657T801 | SM(d18:1/16:0) | neg | 20.40 ± 8.01 |
| M703.5755T801 | SM(d18:1/16:0) | pos | 17.18 ± 5.86 |
| M760.5858T875 | PC(16:0_18:1) | pos | 16.51 ± 2.71 |
| M804.5762T874 | PC(16:0_18:1) | neg | 12.15 ± 2.03 |
| M349.2402T395 | Prostaglandin | neg | 9.35 ± 4.73 |
| M745.5502T739 | SM(d18:1/16:1) | neg | 7.42 ± 1.18 |
| M701.5599T739 | SM(d18:1/16:1) | pos | 5.73 ± 1.94 |
| M338.9687T38 | Unknown | neg | 3.91 ± 3.99 |
| M788.6172T929 | PC(18:0_18:1) | pos | 3.50 ± 0.53 |

**Supplementary Table 4.** Summary of the significantly altered lipid metabolites identified by lipidomic analysis in EVs derived from Y201 MSCs in comparison to EVs derived from Y202 MSCs. Values were normalised by internal standard and particle number. Samples were analysed in the positive and negative ion modes of LC-MS/MS. Unpaired t-test. N=3.

| Label | Annotation | polarity | Mean Y201 (±SD) | Mean Y202 (±SD) | FC (Y201/Y202) | p value |
|---|---|---|---|---|---|---|
| M990.7104T965 | LacCer(d18:1/22:0) | neg | 0.05 ± 0.03 | 0.00 ± 0.00 | 10.40 | 0.003 |
| M415.2121T76 | Unknown | pos | 0.11 ± 0.09 | 1.15 ± 0.66 | 0.10 | 0.018 |
| M1158.7855T998 | Unknown | pos | 0.25 ± 0.05 | 0.12 ± 0.04 | 2.06 | 0.025 |
| M547.5205T775 | Unknown | pos | 0.04 ± 0.03 | 0.29 ± 0.17 | 0.15 | 0.026 |
| M352.2822T153 | Unknown | pos | 0.06 ± 0.05 | 0.37 ± 0.25 | 0.15 | 0.034 |
| M1136.8037T998 | Unknown | pos | 0.38 ± 0.07 | 0.19 ± 0.07 | 1.97 | 0.039 |
| M830.5919T877 | PC(18:1_18:1) | neg | 3.79 ± 0.40 | 2.44 ± 0.54 | 1.56 | 0.039 |
| M518.4939T811 | Unknown | pos | 0.02 ± 0.01 | 0.04 ± 0.01 | 0.47 | 0.044 |
